# A plant pathogen effector blocks stepwise assembly of a helper NLR resistosome

**DOI:** 10.1101/2025.07.14.664264

**Authors:** Benjamin A. Seager, Adeline Harant, Mauricio P. Contreras, Liang-Yu Hou, Chih-Hang Wu, Sophien Kamoun, Jogi Madhuprakash

## Abstract

Helper NLRs function as central nodes in plant immune networks. Upon activation, they oligomerize into inflammasome-like resistosomes to initiate immune signaling, yet the dynamics of resistosome assembly remain poorly understood. Here, we show that the virulence effector AVRcap1b from the Irish potato famine pathogen *Phytophthora infestans* suppresses immune activation by directly engaging oligomerization intermediates of the tomato helper NLR SlNRC3. Cryo-EM structures of SlNRC3 in AVRcap1b-bound and unbound states reveal that AVRcap1b bridges multiple protomers, stabilizing a stalled intermediate and preventing formation of a functional resistosome. Leveraging AVRcap1b as a molecular tool, we also capture an additional SlNRC3 resistosome intermediate showing that assembly proceeds in a stepwise manner from dissociated monomers. These findings uncover a previously unrecognized vulnerability in NLR activation and reveal a pathogen strategy that disrupts immune complex assembly. This work advances mechanistic understanding of resistosome formation and uncovers a previously unrecognized facet of pathogen–plant coevolution.

## Introduction

Animals and plants both rely on one of the most extensive and widespread family of immune receptors, the intracellular nucleotide-binding leucine-rich repeat (NLR) proteins, to defend against pathogen infections (*1-4*). In plants, NLRs are key components of the immune surveillance system recognizing pathogen-secreted molecules known as effectors either directly or indirectly (*5, 6*). A hallmark of NLR activation is their assembly into higher-order oligomeric structures termed resistosomes in plants (*7, 8*), and inflammasomes and apoptosomes in mammals and bacteria (*9-12*). These activated immune complexes prevent pathogen proliferation by triggering a cascade of signalling responses that culminate in localized cell death known as the hypersensitive response. While some NLRs function independently as ‘singletons,’ where a single genetic unit functions in both sensing and signalling, others operate in sub-functionalized pairs (*5, 13*). In these pairs a sensor NLR detects pathogen signals and cooperates with a helper NLR to initiate immune responses (*14, 15*). As a consequence, plants have evolved sophisticated NLR receptor networks to keep pace with rapidly evolving pathogens. A remarkable example of this adaptation is the NRC (NLRs Required for Cell Death) immune network identified in Asterid plants (*13*). This network can comprise dozens of sensor NLRs (disease resistance proteins) that recognize effectors from diverse pathogens and pests, and relay a signal to downstream helper NLRs, known as NRCs, ensuring a robust immune response.

The assembly of higher-order complexes is critical to the function of NLRs in immunity. This assembly process has been previously described for inflammasomes where the NLR family apoptosis inhibitory protein 2 (NAIP2) drives the progressive assembly of NLR family CARD domain-containing protein 4 (NLRC4) into wheel-like structures (*9, 10*). However similar insights in plants have remained limited. More recently, the plant helper NLR N-required gene 1 (NRG1) was shown to undergo induced oligomerization in a heterotrimeric complex with enhanced disease susceptibility 1 (EDS1) and senescence-associated gene 101 (SAG101); however, precise structural details of the assembly process are still lacking (*16*). In their resting states singleton NLRs like hopZ-activated resistance 1 (ZAR1) exist as monomers (*17*) while plant helper NLRs, like NRC2, are present as autoinhibited homodimers (*18, 19*). Although the primed state of ZAR1 has been resolved, the intermediate steps of its oligomeric assembly have not been described. Also, how NRCs transition from dimers into hexameric resistosomes remains unclear and evidence detailing the intermediate stages of this process is lacking.

Pathogens and pests have evolved diverse molecular strategies to counteract immune pathways. In previous studies, we showed that potato pathogens have convergently evolved effectors to suppress NRC helpers; key nodes within the NRC immune network (*20*). One example is the effector SPRYSEC15 (SS15), secreted by the cyst nematode *Globodera rostochiensis*, which specifically binds to the resting-state homodimers of SlNRC1, NbNRC2 and NbNRC3 to prevent their activation (*20, 21*). An unrelated suppressor AVRcap1b—an RXLR-LWY effector from the oomycete *Phytophthora infestans*—binds the host endosomal sorting complex required for transport (ESCRT) membrane trafficking protein Target of Myb 1-like protein 9a (NbTOL9a) to suppress NbNRC2-and NbNRC3-mediated immunity (*20*). We recently showed that, in addition to binding NbTOL9a, AVRcap1b selectively associates with the activated state of NRCs, but not their resting state homodimers (*22*). Here, we reasoned that AVRcap1b could serve as a molecular probe to investigate the activation mechanisms of helper NLRs and determined the cryo-electron microscopy (cryo-EM) structures of activated tomato SlNRC3 in both AVRcap1b-bound and unbound states. These structures reveal how AVRcap1b suppresses NRC activation and, importantly, capture discrete assembly intermediates, providing new insights into the dynamics of resistosome formation.

## Results

### AVRcap1b binds activated SlNRC3 and alters resistosome formation

We previously showed that AVRcap1b suppresses immune cell death by bridging the membrane trafficking protein NbTOL9a to activated NbNRC2 (*22*). However, repeated attempts to purify an NbNRC2–AVRcap1b complex remained unsuccessful. This prompted us to investigate the spectrum of AVRcap1b suppression activity across a wide phylogenetic spectrum of Solanaceae species. To this end, we assayed AVRcap1b suppression of NRC1, NRC2, and NRC3 orthologs from *Nicotiana benthamiana, Capsicum annuum* (pepper), *Solanum tuberosum* (potato), and *Solanum lycopersicum* (tomato). We used the well-established NRC activation system based on the coat protein (CP) of *Potato virus X* (PVX) and its matching sensor NLR, the potato disease resistance protein Rx (**Fig. 1A**). While AVRcap1b exhibited variable suppression of NRC1 and NRC2 orthologs it consistently and fully suppressed the hypersensitive cell death mediated by all four NRC3 orthologs (**Fig. 1B & fig. S1A, B**). This broad-spectrum suppression points to a potentially robust interaction between effector and NLR, leading us to pivot to NRC3 to investigate the biochemical mechanisms of AVRcap1b-mediated immune suppression.

**Fig. 1:**
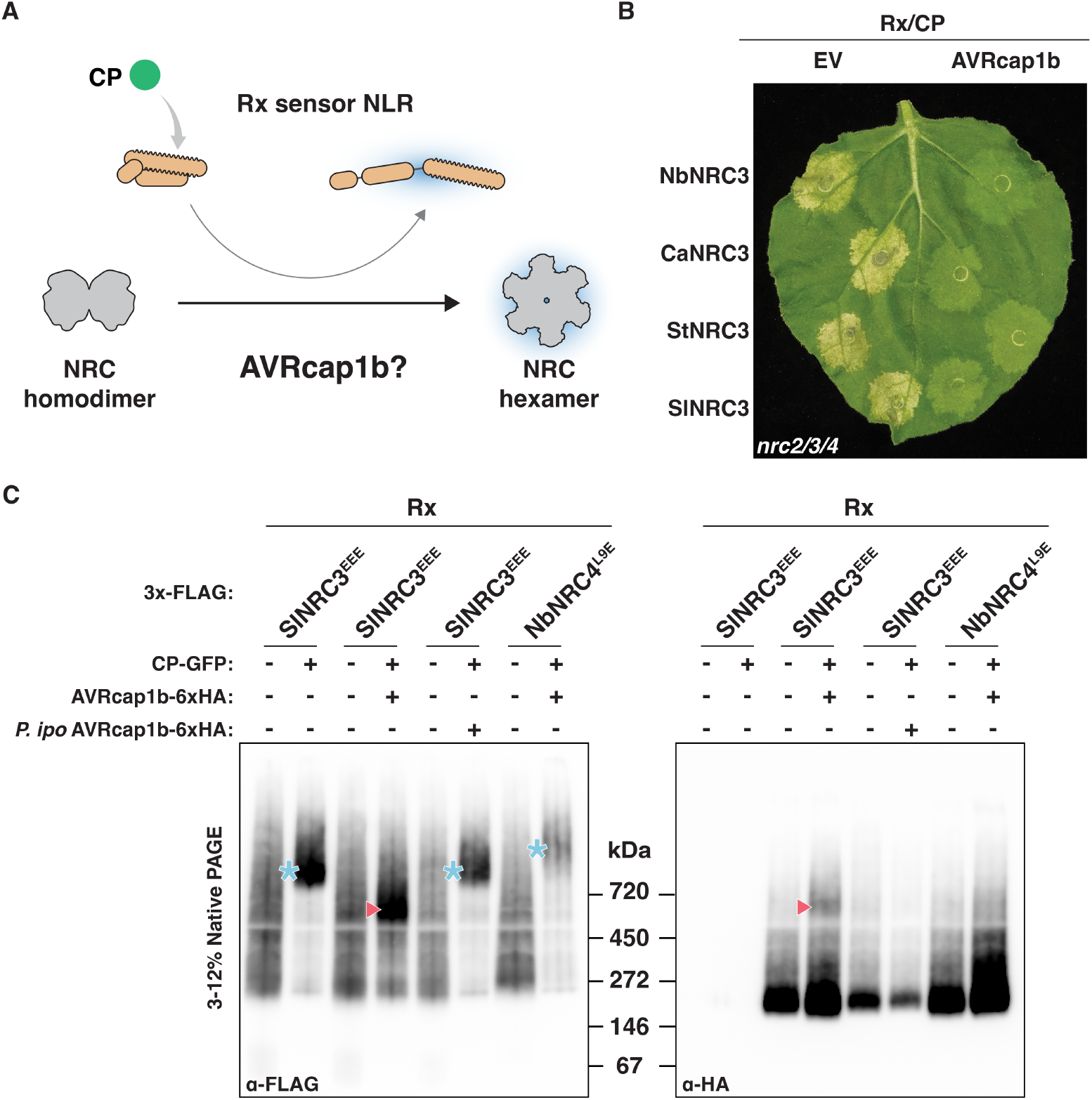
AVRcap1b suppresses NRC3-mediated plant immunity by blocking resistosome assembly. A) Schematic showing the well-characterized sensor Rx (orange)/ *Potato virus X* (PVX) coat protein (CP (green) activation of NRCs used to assess AVRcap1b suppression B) Representative leaf image from *N. benthamiana nrc2/3/4* knockout plants showing hypersensitive cell death response after co-expression of either an empty vector (EV) or AVRcap1b with Rx, PVX-CP, and different NRC3 orthologs. (Nb: *Nicotiana benthamiana*, Ca: *Capsicum annuum*, St: *Solanum tuberosum*, Sl: *Solanum lycopersicum*) C) BN-PAGE analysis showing inactive and activated Rx co-expressed with either SlNRC3^EEE^ or NbNRC4^L9E^, in the presence or absence of AVRcap1b. Sensor-helper combinations were co-infiltrated along with GFP or PVX-CP-GFP. *Phytophthora ipomoeae* (*P. ipo*) AVRcap1b, which does not suppress NRC3 activity, was used as a negative control. Total protein extracts were immunoblotted with the indicated antibodies (bottom left). Corresponding SDS-PAGE blots are presented in **fig. S1C**. Approximate molecular weights (in kilodaltons) are shown between the two blots. Blue asterisks denote higher molecular weight complexes (hexameric resistosomes) and red triangles denote a high-order oligomeric complex of SlNRC3^EEE^ and AVRcap1b that is intermediate in size between SlNRC3 homodimers and hexamers. The experiment was repeated three times with similar results.

Next, we used blue-native polyacrylamide gel electrophoresis (BN-PAGE) assays to examine the degree to which AVRcap1b affects SlNRC3 oligomerization following PVX-CP and Rx activation **(Fig. 1C & fig. S1C)**. As in previous work, we used mutants in the MADA motif of the N-terminal α1 helices—which assemble into a funnel-like membrane pore in the resistosome—because they abolish cell death induction without compromising receptor activation or oligomerization (*23, 24*). In the presence of AVRcap1b and upon activation with PVX-CP and Rx, a new band consisting of a high-order oligomeric complex of SlNRC3 and AVRcap1b appeared that was distinct from the larger hexameric resistosome of SlNRC3 **(Fig. 1C & fig. S1C)**. In addition, AVRcap1b reduced the intensity of the hexameric oligomer of activated SlNRC3, suggesting that it interferes with resistosome assembly. For negative controls, we found that the AVRcap1b ortholog from *Phytophthora ipomoeae*, which doesn’t suppress NRC activity (*22*), didn’t affect SlNRC3 oligomerization and AVRcap1b didn’t affect oligomerization of NbNRC4 as previously reported (*20*). We conclude that AVRcap1b likely interferes with resistosome assembly by stabilizing an intermediate state.

We previously showed that the AVRcap1b^P92E^ mutant is compromised in NbTOL9a binding and NRC suppression (*22*). Given our finding that AVRcap1b also binds activated SlNRC3, we investigated the degree to which AVRcap1b^P92E^ retains any suppression activity. To this end, we performed two independent experiments. First, we took advantage of the recently developed copper inducible transcriptional activation system to perform a time course assay of SlNRC3 hypersensitive cell death following co-expression of PVX-CP and SlNRC3 followed by copper-induction of Rx one day later (**fig. S2A**)(*25*). In these assays, the P92E mutant of AVRcap1b retained partial suppression of SlNRC3-mediated cell death when compared to the wild-type AVRcap1b and the mock negative control or the *P. ipomoeae* AVRcap1b, which binds NbTOL9a but doesn’t suppress NRCs (**fig. S2**)(*22*). In independent time course assays, in which the PVX-CP, Rx and SlNRC3 were co-expressed and the cell death response was monitored over time, the AVRcap1b^P92E^ mutant also showed partial suppression of SlNRC3, notably in the early days after activation (**fig. S2B**). These findings independently corroborate our earlier observations that AVRcap1b can interfere with SlNRC3 resistosome formation, and reveal that SlNRC3 retains, at least in part, an NbTOL9a-independent suppression activity.

### Cryo-EM analysis reveals an AVRcap1b-bound SlNRC3 resistosome intermediate

Our initial experiments prompted us to resolve the structure of the AVRcap1b-SlNRC3 complex using cryo-EM. We purified SlNRC3 activated by PVX-CP and sensor Rx in the presence and absence of AVRcap1b following transient expression in *N. benthamiana* and used cryo-EM to determine the structure of both complexes to understand how the effector could be interfering with immune activation. Activated SlNRC3 formed a hexameric resistosome that resembled the previously reported structures of activated NbNRC2 and NbNRC4 (**Fig. 2A, fig. S3-S5 & Table S1**)(*8, 26*). Intriguingly, activated SlNRC3 in complex with AVRcap1b showed clear density upon two-dimensional (2D) classification which upon three-dimensional reconstruction revealed a structure comprising three activated SlNRC3 protomers bound by the L-shaped AVRcap1b effector **(Fig. 2B-D, fig. S6-S9 & Table S1)**. This structure was consistent with the reduced size of the band observed in the BN-PAGE (**Fig. 1)** confirming that AVRcap1b can disrupt assembly of the hexameric resistosome. AVRcap1b interacts with all three SlNRC3 protomers on the interface that, in a fully formed resistosome, would normally be masked by the second half of the hexamer. The effector wraps around the bottom of the three protomers making contact with the underside of the assembled NLRs. Structural superimposition of the hexameric resistosome and the SlNRC3-AVRcap1b intermediate complex confirms that the latter corresponds to one half of the complete hexameric resistosome (**Fig. 2E**), with minimal difference in root mean square deviation (RMSD = 0.793 Å) (**fig. S10**).

**Fig. 2:**
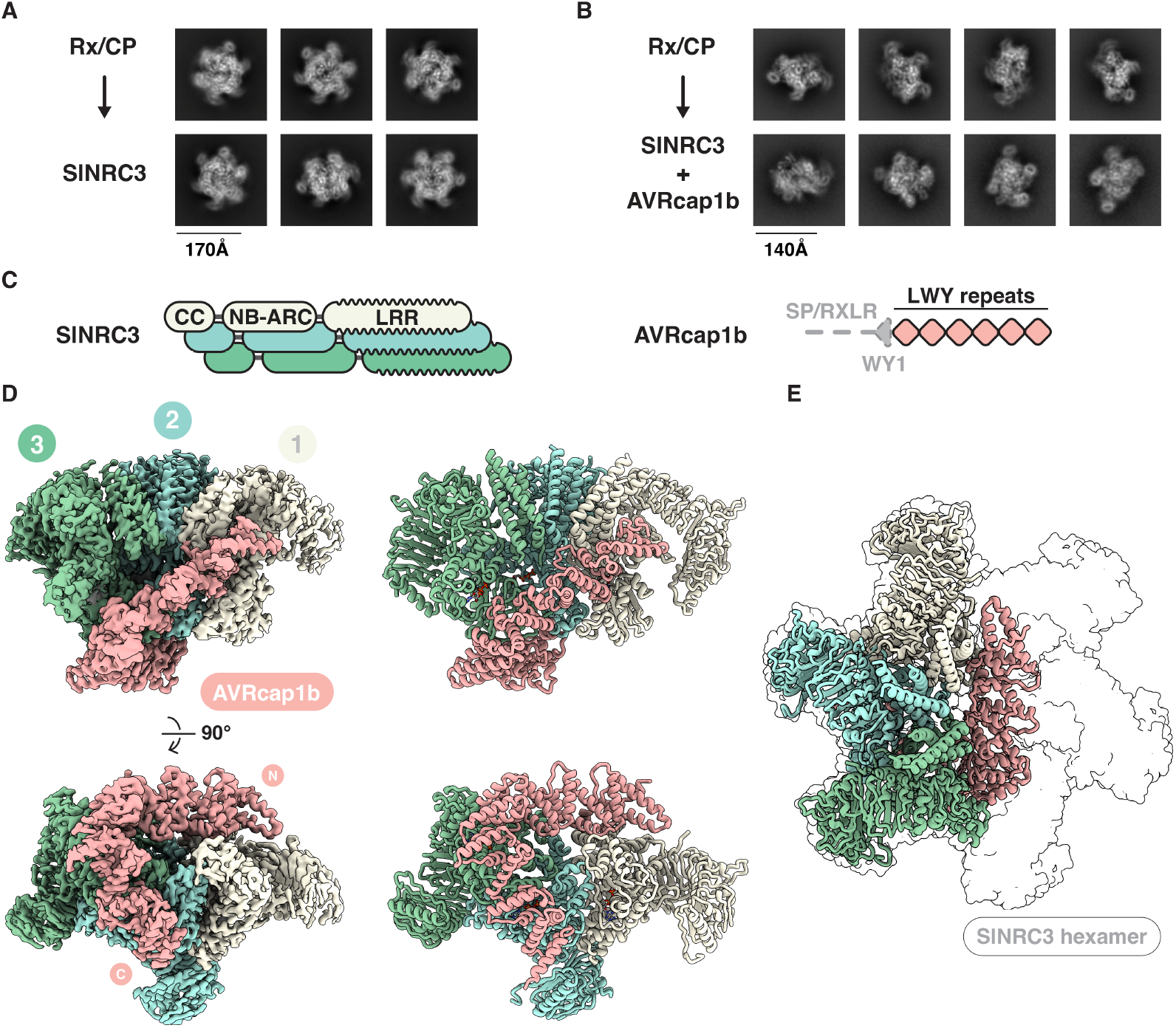
Cryo-EM structure reveals an intermediate form of the SlNRC3 resistosome bound to AVRcap1b. A & B) Representative cryo-EM 2D class averages showing distinct views of the SlNRC3 resistosome and its intermediate assembled in the presence of AVRcap1b. C) Schematic representation of the domain organization of SlNRC3 (CC: coiled-coil, NB-ARC: nucleotide binding adaptor shared by APAF-1, plant R proteins, and CED-4, LRR: leucine rich repeat) the effector AVRcap1b (SP/RXLR: signal peptide and RXLR motif) D) Composite cryo-EM density map (left) and the corresponding atomic model (right) of the SlNRC3-AVRcap1b complex, shown in two orientations. SlNRC3 protomers are shown in white, cyan and green, and numbered according to the order in which they contact AVRcap1b from N- to C- terminus. AVRcap1b is depicted in light pink, with annotated N- and C-termini. Notably, the cryo-EM reconstruction lacks interpretable density for the N-terminal WY1 domain of AVRcap1b, suggesting conformational flexibility. E) Superimposition of the SlNRC3-AVRcap1b structure onto the SlNRC3 hexamer reveals that AVRcap1b sterically hinders complete resistosome formation, stabilizing a three- protomer assembly intermediate.

### AVRcap1b forms an extensive interface with the SlNRC3 assembly intermediate

The L-shaped AVRcap1b effector binds to the three-protomer intermediate through an extensive interface involving the coiled-coil (CC), winged helix domain (WHD) and nucleotide binding (NB) domains (**Fig. 3A & Table S2**). The WY1 domain of AVRcap1b, known to contribute to the NbTOL9a binding (*20*), was not resolved in our structure suggesting flexibility of this domain that may be important for NbTOL9a engagement. The remaining LWY domains (LWY2-7) collectively form the interface between AVRcap1b and the three SlNRC3 protomers **(Fig. 3B)**. LWY2 primarily engages with the CC domain of protomer-1 but also makes minor contact with the WHD. LWY3 makes some minor contacts with the CC domains of protomers 2 and 3. Notably, a loop within LWY3 interacts with Leu134 of all three protomers, a residue that contributes to the SlNRC3 hexamer pore **(Fig. 3C)**. LWY4 and LWY5 establish minimal contacts with the NB domain of protomer-3 where AVRcap1b bends to form contacts with the underside of both protomers 2 and 3. Among the LWY domains, LWY6 contributes to the most extensive interaction interface, with a buried surface area of 729 Å^2^, through multiple hydrogen bonds and hydrophobic interactions. Specifically, Asp566 of LWY6 forms a salt bridge with Lys256 of SlNRC3 **(Fig. 3D)**. Furthermore, Arg513 of LWY6 forms hydrogen bonds with the backbone of residues Glu203 and Tyr204 of protomer-3, while Phe207 of SlNRC3 is captured within a hydrophobic pocket of LWY6. This same region of SlNRC3 also forms part of the binding site for LWY7 on protomer-2. In this interaction, Tyr204 is buried within a hydrophobic pocket of LWY7, where it forms a hydrogen bond with Asp609 **(Fig. 3E)**. Another key residue, Glu205 of protomer-2, establishes a salt bridge with Arg652 of AVRcap1b. Phe207, which plays a crucial role in the LWY6 interaction with protomer-3, also contributes significant buried surface area to the LWY7 interface with protomer-2. Together, LWY6 and LWY7 account for 61% (1274 Å^2^ out of a total of 2088 Å^2^) of the total buried surface area between AVRcap1b and the three SlNRC3 protomers, underscoring their key role in effector binding.

**Fig. 3:**
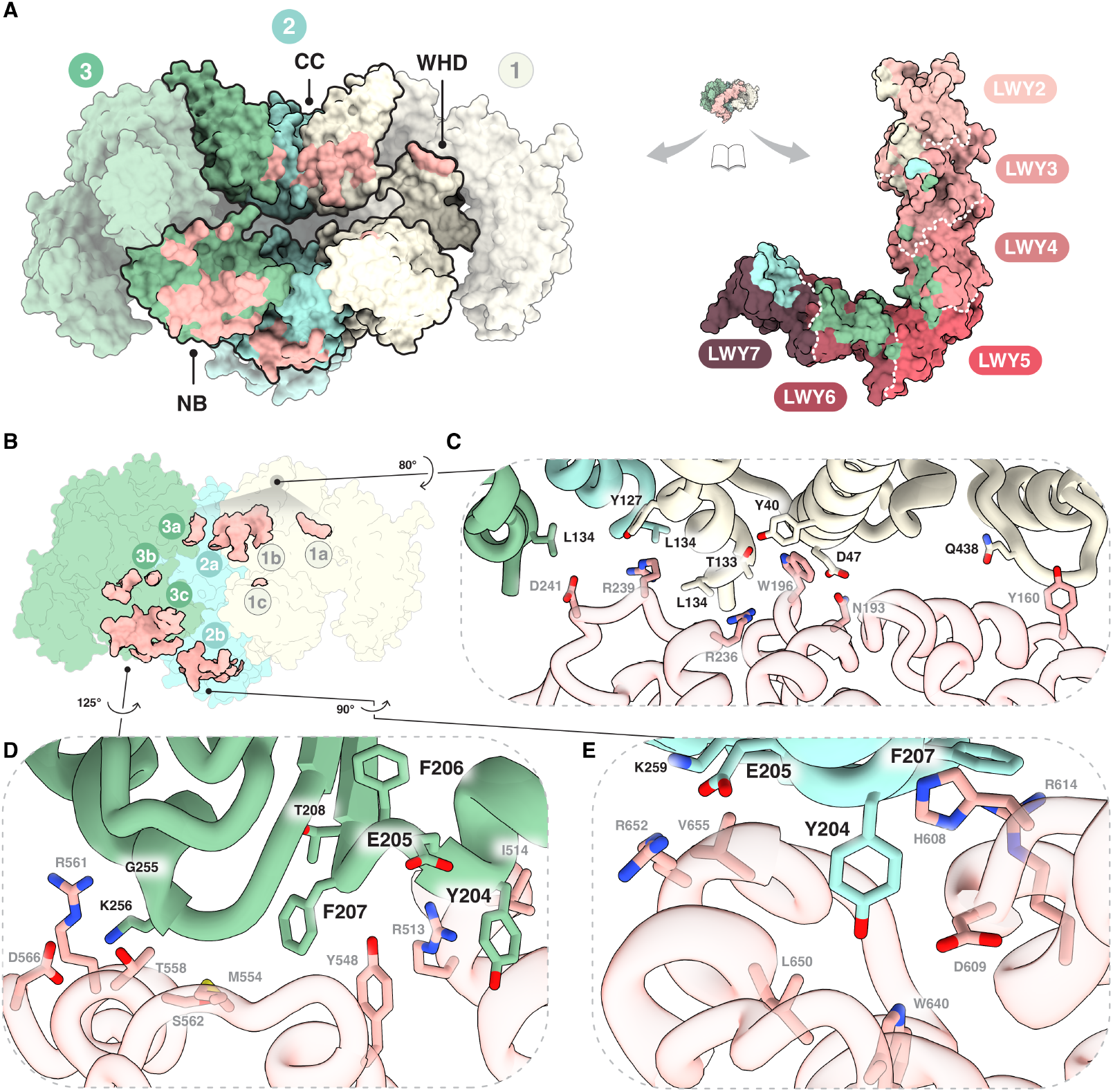
AVRcap1b binds across all three SlNRC3 protomers through multiple interfacial contacts. A) Open-book view illustrates the complementary interaction surfaces between SlNRC3 (left) and AVRcap1b (right). Surface representation of the SlNRC3-AVRcap1b complex, showing individual SlNRC3 protomers in distinct colors, with the domains (CC, NB, WHD) involved in contacts highlighted. AVRcap1b is displayed in a domain-colored scheme, highlighting the boundaries between the LWY domains. B) Mapping of the interfacial surfaces across all three SlNRC3 protomers revealed multiple contact points (1a–1c, 2a–2b, 3a–3c) with distinct LWY domains of AVRcap1b. C–E) Close-up views of selected interface regions highlight key residues involved in stabilizing the complex. Contacting residues from SlNRC3 and AVRcap1b are shown in stick representation. Conserved SlNRC3 residues, Y204, E205, and F207, form a crucial contact patch that recurs specifically across LWY6 and LWY7 interactions. Together, these distributed binding interfaces stabilize the AVRcap1b bound SlNRC3 intermediate and prevent resistosome assembly.

### N-terminal truncations of AVRcap1b reveals the stepwise assembly of SlNRC3 resistosome

The ability of AVRcap1b to interfere with SlNRC3 resistosome assembly prompted us to exploit AVRcap1b as a tool to investigate the assembly process. We hypothesized that removing one or more N-terminal LWY domains would facilitate the incorporation of additional activated SlNRC3 protomers based on structural comparisons between the hexameric resistosome and the AVRcap1b-bound assembly intermediate. This, in turn, could capture resistosome assembly intermediates with varying stoichiometry. To this end, we generated three N-terminal truncations of AVRcap1b given that LWY6 and LWY7 were critical for NRC binding. We co-expressed these truncated variants with PVX-CP and Rx activated SlNRC3 in *N. benthamiana*, purified the complexes, and assessed their ability to block resistosome assembly using negative stain electron microscopy. Remarkably, negative-stain analysis of the AVRcap1b truncation containing only LWY3-7 revealed an assembly intermediate consisting of four SlNRC3 protomers (**Fig. 4**). The two other truncations that were tested did not suppress cell death, and did not co-purify with SlNRC3, suggesting complete loss of binding and were therefore omitted from negative stain analysis (**fig. S11**). The successful incorporation of an additional protomer with a truncated AVRcap1b variant is consistent with our structural analysis and suggests that resistosome assembly proceeds in a stepwise manner through the progressive addition of activated SlNRC3 protomers culminating in a functional hexameric resistosome.

**Fig. 4:**
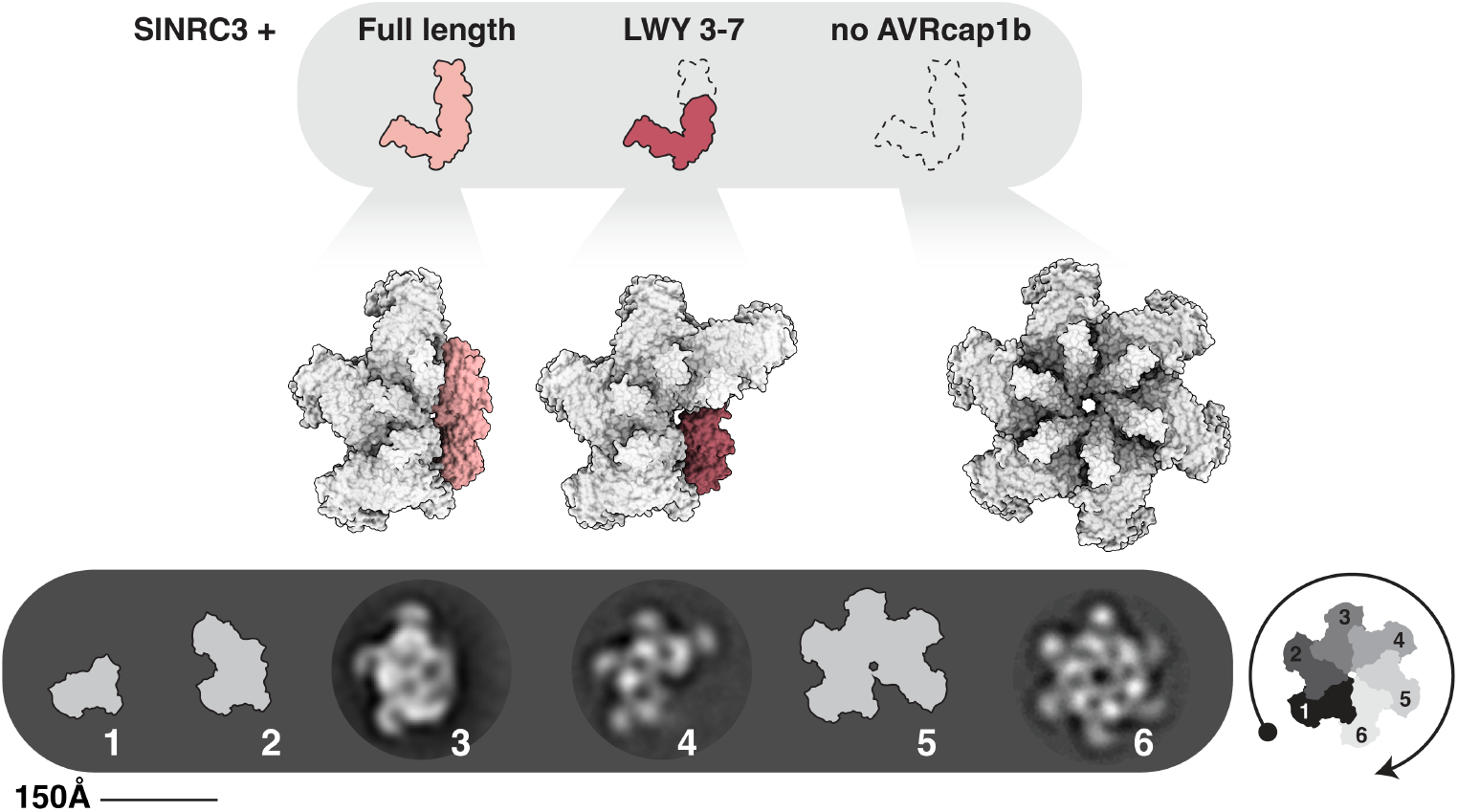
AVRcap1b disrupts stepwise assembly of the SlNRC3 resistosome. *Top*: Schematic representation of full-length AVRcap1b and its N-terminal truncation (LWY3–7), both of which bind SlNRC3; dashed outlines indicate absence of AVRcap1b domains. *Middle*: Structural models of SlNRC3 oligomerization states in the presence of full-length AVRcap1b, AVRcap1b^LWY3–7^, or no effector. The central model, representing an intermediate stalled by AVRcap1b^LWY3–7^, was generated using AlphaFold 3 (*27*). *Bottom*: Negative-stain 2D class averages suggest a progression of oligomeric intermediates (with inferred states for species 1, 2, and 5 in cartoon representation), culminating in a complete hexamer in the absence of AVRcap1b. Full-length AVRcap1b and AVRcap1b^LWY3–7^ stall SlNRC3 at distinct sub- hexameric states, indicating that AVRcap1b interferes with resistosome formation by trapping assembly intermediates.

### Mutational analysis validates the SlNRC3-AVRcap1b complex interface

Structural analysis of the SlNRC3-AVRcap1b complex revealed that SlNRC3 Tyr204, Glu205, Phe206, and Phe207 (hereafter, the YEFF motif) are simultaneously engaged by LWY6 and LWY7 of AVRcap1b. This YEFF sequence resides at the periphery of the NB domain and forms a conserved motif across NRC3 orthologs from all examined Solanaceae species (**fig. S12**).

To determine the degree to which the YEFF motif contributes to the SlNRC3-AVRcap1b interface, we performed mutagenesis experiments on SlNRC3. Wild-type tomato NRC3 (SlNRC3^WT^) is autoactive in *N. benthamiana* making mutational assays challenging (**Fig. 5A**). Therefore, we initially performed hypersensitive cell death assays by mutating equivalent residues in the *N. benthamiana* ortholog NbNRC3. Out of ten different mutations tested, including single and double mutants, NbNRC3^E207A^ and NbNRC3^F209A^, along with three double mutants (NbNRC3^Y206A/E207A^, NbNRC3^Y206A/F209A^, and NbNRC3^E207A/F209A^) became insensitive to AVRcap1b suppression (**Fig. 5A & fig. S13**). However, with the exception of NbNRC3^Y206A/E207A^, the remaining four mutants displayed a sensitized ‘trigger-happy’ phenotype (*28*) where cell death was observed when co-expressed with the sensor Rx alone. Additional mutations targeting residues that contact LWY6 (NbNRC3^R211A^, NbNRC3^K258A^, and NbNRC3^R211A/K258A^) did not markedly affect suppression by AVRcap1b, suggesting that the interactions formed by LWY7 are more critical for effector binding (**fig. S13B**).

**Fig. 5:**
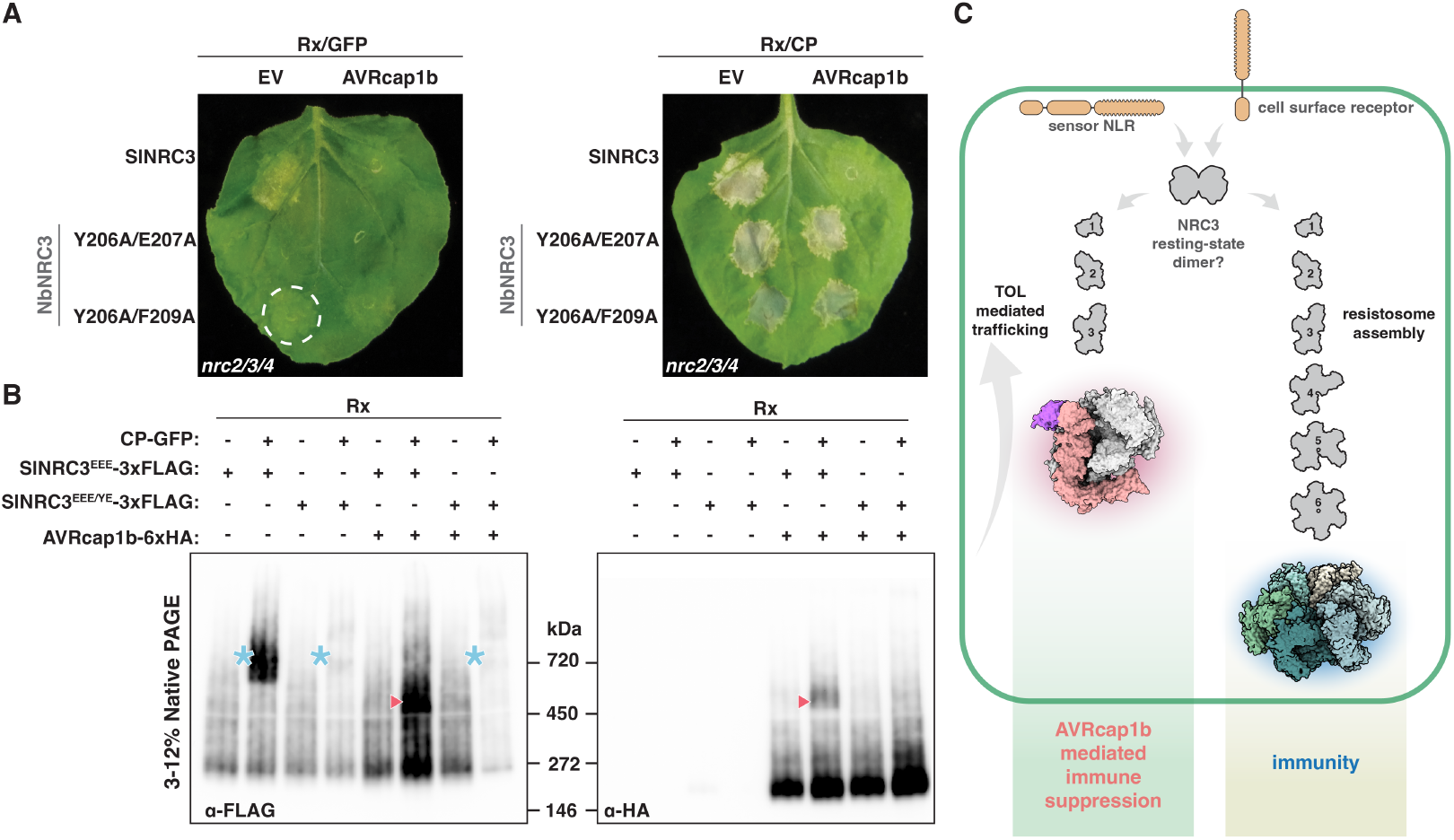
Mutations in the YEFF motif of SlNRC3 disrupt AVRcap1b-mediated suppression. A) Representative *N. benthamiana nrc2/3/4* knockout leaf showing hypersensitive cell death response following co-expression of PVX-CP, Rx sensor, AVRcap1b, and SlNRC3 or its double mutants. The Y206A/E207A mutant restored immune activity without inducing autoactivity, whereas the mutant Y206A/F209A (highlighted with a white circle) displayed mild autoactivity. B) BN-PAGE analysis of inactive and activated Rx co-expressed with either SlNRC3^EEE^ or SlNRC3^EEE/YE^, in the presence or absence of AVRcap1b. Sensor-helper combinations were co-infiltrated along with GFP or PVX-CP-GFP. Total protein extracts were immunoblotted with the indicated antibodies (bottom left corner). Corresponding SDS-PAGE blots are presented in **fig. S17**. Approximate molecular weights (in kilodaltons) are indicated between the two blots. Blue asterisks denote higher molecular weight complexes (hexameric resistosomes) and red triangles denote a shift in the higher-order oligomeric state of SlNRC3^EEE^ in the presence of AVRcap1b. The same band is detected by α-HA, indicating that AVRcap1b co-migrates with an intermediate SlNRC3^EEE^ resistosome complex. In contrast, SlNRC3^EEE/YE^ assembled into high-molecular-weight complexes regardless of AVRcap1b, suggesting that these substitutions do not impair resistosome formation but prevent effector binding. *Note*: The SlNRC3^EEE/YE^ corresponds to the Y204A/E205A mutant introduced into the N-terminal MADA-mutated SlNRC3 background to prevent cell death induction upon activation. The experiment was repeated three times with similar results. C) Proposed working model of immune activation and suppression. Upon effector or PAMP recognition by intracellular sensors or cell-surface receptors, respectively, resting-state NRC3 dissociates into monomers and assembles into resistosomes in a stepwise manner. The fully assembled complex inserts into the plasma membrane to trigger localized cell death and limit pathogen spread. AVRcap1b binds to NRC3 assembly intermediates, halting resistosome formation and suppressing immune activation. The NbTOL9a and NRC3 interfaces of AVRcap1b are mutually exclusive (**fig. S18**), allowing the stalled complexes to be potentially trafficked via the TOL-ESCRT membrane remodeling pathway. Numbers indicate protomer count within each oligomeric complex.

To further validate these findings, we mutated residues in LWY7 of AVRcap1b determined from the cryo-EM structure to contact the NB domain of SlNRC3. Four single and two double AVRcap1b mutants in LWY7 lost the ability to suppress SlNRC3, underscoring the crucial role of the ‘LWY7–NB’ interface and orthogonally validating the structural model **(fig. S14)**. In a complementary experiment, the *P. ipomoeae* AVRcap1b ortholog, which doesn’t suppress NRC3, gained NRC3 suppression activity upon swapping of its LWY7 domain with that from *P. infestans* AVRcap1b, further demonstrating the critical role of this domain in engaging with the activated NLR (**fig. S15**).

One mutant in the YEFF motif that became insensitive to AVRcap1b suppression, NbNRC3^Y206A/E207A^, was neither autoactive nor trigger-happy (**Fig. 5A & fig. S13**). To further determine the importance of this motif, we introduced Y206A/E207A in the corresponding residues of the NRC3 orthologs described above: CaNRC3, StNRC3, and SlNRC3 (**Fig. 1**), generating CaNRC3^Y204A/E205A^, StNRC3^Y204A/E205A^, and SlNRC3^Y204A/E205A^. Besides the previously tested NbNRC3^Y206A/E207A^, two of these NRC3 double mutants StNRC3^Y204A/E205A^ and SlNRC3^Y204A/E205A^ became insensitive to suppression by AVRcap1b, whereas CaNRC3^Y204A/E205A^ remained partially suppressed (**fig. S16**).

We further investigated the impact of the YEFF motif on AVRcap1b suppression of SlNRC3 using BN-PAGE by assaying the Y204A/E205A mutations in the presence or absence of the effector (**Fig. 5B & fig. S17**). Upon activation with PVX-CP and Rx, the mutant SlNRC3^Y204A/E205A^ assembled into high-molecular-weight resistosome complexes, although the bands weren’t as prominent as with the SlNRC3 control (**Fig. 5B**). However, in the presence of AVRcap1b, SlNRC3^Y204A/E205A^ didn’t associate with AVRcap1b in an intermediate band (presumably the 3:1 SlNRC3: AVRcap1b complex) as clearly observed with SlNRC3 (**Fig. 5B**). These results independently confirm that the Y204A/E205A substitutions disrupt AVRcap1b binding without compromising SlNRC3 resistosome assembly and provides further support for a model in which the effector targets a conserved NB-domain interface to stall resistosome assembly of a helper NLR at a sub-hexameric state (**Fig. 5C**).

## Discussion

Our understanding of the structural basis of plant immune receptors primarily stems from studies on how they recognize pathogen effectors to activate immunity (*7, 29, 30*). Here, we reveal a novel aspect of pathogen–plant coevolution through the cryo-EM structure of a pathogen immunosuppressor in complex with an activated helper NLR trapped in a sub-hexameric state. We captured the effector AVRcap1b from the Irish potato famine pathogen *P. infestans* in complex with stalled three-protomer intermediate of the tomato helper NLR SlNRC3. This, in addition to our findings using truncated AVRcap1b variants, suggest that helper NRCs probably undergo activation from primed monomers dissociated from the autoinhibited homodimeric resting state (*18, 19*) through stepwise oligomerization into a homohexameric resistosome (*8*) (**Fig. 5C**). These structural snapshots of intermediate states that arise during resistosome formation complement previously reported static structures of resting and resistosome endpoint states. This sheds new light on how NLRs dynamically shift from resting to activated states, adding to limited studies on mammalian NLR oligomerization (*9, 10*).

Previous studies have established that paired NLRs like NLRC4 and NRG1 oligomerize through interactions with heterodimeric (NLRC4-NAIP2) or heterotrimeric (NRG1-EDS1-SAG101) complexes (*9, 10, 16*). In contrast, oligomerization of NRC helpers doesn’t involve additional partner proteins. Our findings support the activation-and-release model for sensor-helper pairs in the NRC network (*24*), where effector recognition induces conformational changes in sensor NLRs, such as Rx, that transiently engage and activate downstream NRC helpers. Given that the protein complexes we analyzed, whether by BN-PAGE or *in planta* purification, did not capture the Rx sensor, we conclude that any interactions between Rx and SlNRC3 are transient. The helper NLRs dissociate to assemble the resistosome independently, with no evidence of sensor-mediated nucleation.

There are only few examples of pathogen or parasite effectors that directly target NLRs to inhibit their activities. The potato cyst nematode effector SS15 binds the NB domain of autoinhibited NRCs, physically preventing conformational rearrangements required for activation (*20, 21, 31*). In contrast, the oomycete effector AVRcap1b, which convergently evolved to also suppress NRC2 and NRC3, interferes with resistosome assembly exposing a previously unrecognized vulnerability in NLR activation. Both molecular mechanisms of helper NLR inhibition must be effective, but they may be linked to physiological constraints. In the case of SS15, the stoichiometry of the effector versus the NLR must be favorable to the pathogen to ensure effective immunosuppression. Also, the suppressor needs to be delivered before the activating nematode effector given that SS15 cannot act on the activated NRC state. In the case of AVRcap1b, timing constraints on its delivery may be more relaxed, as it acts downstream of NRC activation. In addition, AVRcap1b direct inhibition of NRC oligomerization only results in partial immunosuppression and depends on the host membrane trafficking protein NbTOL9a for full suppression activity (**fig. S2**) (*20, 22*).

Therefore, AVRcap1b doesn’t function as a *bona fide* NLR inhibitor like SS15 but rather bridges activated NLRs to NbTOL9a for full immunosuppression. Contrasting mechanisms of the two effectors from widely divergent parasites (metazoan vs. oomycete) that have convergently evolved to suppress critical NRC nodes in an immune receptor network underscore how different points of attack can carry different trade-offs for both pathogen and host.

AVRcap1b adopts an original L-shaped structure distinct from other previously studied RXLR-LWY effectors in *Phytophthora*, confirming its structural divergence within this family (*22*). It acts as a molecular bridge, using its WY1–LWY2 domains to bind NbTOL9a at a site distinct from the NRC interface (**Fig. 5C & fig. S18**). This dual-targeting mechanism, which links immune suppression to membrane trafficking, echoes similar strategies seen in effectors like the aster yellows phytoplasma effector SAP05 and the *Phytophthora sojae* RXLR-LWY effector PSR2 (*32, 33*). However, unlike PSR2, which modulates a single regulatory hub, AVRcap1b coordinates two distinct host pathways, dragging NRC resistosome assembly intermediates to the NbTOL9a ESCRT membrane trafficking pathway (*22*). This functional and evolutionary modularity highlights how AVRcap1b, like other modular RXLR-LWY effectors (*33*), expands effector versatility by bridging across distinct pathways.

Our cryo-EM structure revealed how the L-shape of AVRcap1b forms a complementary shape that accommodates the resistosome assembly intermediate of SlNRC3. Analysis of conservation across homologous RXLR-LWY effector sequences from *Phytophthora* shows that the NRC binding regions exhibit the lowest sequence conservation of surface exposed residues (**fig. S19)** (*22*). While AVRcap1b from *P. infestans* specifically targets a subset of NRCs, other *Phytophthora* species have presumably evolved AVRcap1b-like effectors to target a unique set of host proteins. Whether other L-shaped RXLR-LWY effectors of the AVRcap1b family also inhibit NLRs, and the extent to which diversification of the SlNRC3 binding interface of AVRcap1b reflects adaptations to suppress a broad range of immune receptors, is an intriguing hypothesis that deserves to be explored.

## Acknowledgments

The authors thank J. Win, J. Kourelis (The Sainsbury Laboratory) and M. Webster (John Innes Centre) for their helpful suggestions. We thank J. Richardson (John Innes Centre) for aiding in negative stain data acquisition and N. Lukoyanova and S. Chen (Birkbeck University of London) for cryo-EM data acquisition. We thank D. Lawson (John Innes Centre) for assistance with electron microscopy data storage and computing resources. We thank the Horticultural Services and Scientific Support teams (John Innes Centre and The Sainsbury Laboratory) for their assistance. J.M. acknowledges the support from University of Hyderabad, India.

## Author contributions

Conceptualization: B.A.S., J. M., S.K., Methodology: B.A.S., A.H. M.P.C., L.-Y. H, C.-H. W., J.M. Investigation: B.A.S, M.P.C., L.-Y.H., J.M. Formal analysis: B.A.S, M.P.C., L.-Y.H., C.-H. W., J.M. Resources: B.A.S., A.H. M.P.C., J.M., Writing – original draft: B.A.S., J.M. Writing – review & editing: B.A.S., A.H. M.P.C., L.-Y. H, C.-H. W., S.K., J.M. Visualization: B.A.S. M.P.C., L.-Y. H., J.M. Supervision: S.K., J.M. Project administration: S.K., J.M. Funding acquisition: S.K.

## Data and materials availability

All maps and models have been deposited in PDB and EMDB with the following accessions: SlNRC3 hexamer (PDB: 9RI9, EMDB: EMD-53990), SlNRC3-AVRcap1b complex (PDB: 9RIA, EMDB: EMD-53991 for composite map, EMD-53988 for consensus reconstruction, and EMD-53989 for focused refinement of AVRcap1b). Constructs used in this study will be available upon request subject to material transfer agreements (MTA).

## Materials and Methods

### Plant growth conditions

CRISPR-engineered *N. benthamiana nrc2/3/4* knockout lines were cultivated in a controlled-environment growth chamber maintained at 22–25 °C, with relative humidity between 45% and 65%, under a 16-hour light / 8-hour dark photoperiod.

### Plasmid constructions

Constructs were generated using the Golden Gate Modular Cloning (MoClo) system (*34*), along with the MoClo Plant Parts Kit (*35*), unless otherwise specified. The expression vectors for *PVX-CP-eGFP, Rx*-2xV5, and *AVRcap1b*-6xHA, used in protein purification assays, were previously described (*22*). For this study, *SlNRC3^EEE^*-3xFLAG, *NbNRC3^EEE^*-3xFLAG, and corresponding mutant variants were cloned into the binary vector pJK001c, incorporating a dual 35S + Ω promoter (pICH51288), a 35S terminator (pICH41414), and a C-terminal 3xFLAG tag (pICSL50007) (*36*). Truncated forms of *AVRcap1b, AVRcap1b^LWY3–7^, AVRcap1b^LWY4–7^*, and *AVRcap1b^LWY5–7^*, were cloned into the binary vector pICSL47742, containing a long 35S + Ω promoter (pICH51266), an *ocs* (octopine synthase) terminator (pICH41432), and a C-terminal 6xHA tag (pICSL50009). Cloning strategies and sequence validation were performed using Geneious Prime (version 2021.2.2; https://www.geneious.com).

### Cell death assays

Effector proteins and NLR immune receptors were transiently expressed in *N. benthamiana nrc2/3/4* knockout leaves using *Agrobacterium tumefaciens*–mediated infiltration, as previously described (*37*). Briefly, 4–5-week-old plants were infiltrated with suspensions of *A. tumefaciens* strain GV3101 pMP90 harboring the respective expression constructs. Bacterial cultures were resuspended in infiltration buffer (10 mM MES, 10 mM MgCl_2_, 150 μM acetosyringone, pH 5.6) and adjusted to the following final optical densities (OD_600_): 0.3 for NRCs and their variants, 0.1 for Rx, 0.1 for PVX-CP, and 0.3 for AVRcap1b. The total OD_600_ for all combined suspensions was maintained at 0.8. Cell death was assessed 5 days post agroinfiltration. All hypersensitive cell death assays were scored on a modified 0-7 scale and quantification was carried out in Rstudio.

### Copper inducible time-course cell death assay

To determine the dynamics of cell death, we performed a time-course cell death assay using the copper-inducible system (*38*). Leaves from *N. benthamiana nrc2/3/4* knockout plants were agroinfiltrated with CBS4::HyP5sm-Rx, 35S::CP, 35S::SlNRC3, 35S::CUP2-p65, and CBS4::OSL5, in combination with one of the following constructs: AVRCap1b, AVRCap1b^P92E^, *P. ipomoeae* AVRCap1b, or an empty vector (EV) control. Bacterial suspensions were adjusted to the following final OD_600_ values: 0.1 for CBS4::HyP5sm-Rx, 35S::CUP2-p65, CBS4::OSL5, and each AVRCap1b variant; and 0.2 for 35S::CP and 35S::SlNRC3. The total OD_600_ of each mixed suspension was brought to 0.8 using an EV suspension. At one day post-agroinfiltration, 10 μM copper sulfate solution was infiltrated into the leaves to induce hypersensitive cell death. Subsequently, leaves were detached and placed on 1% agarose gel to prevent dehydration and imaged using the PerkinElmer IVIS Lumina III imaging system. Autofluorescence was excited at 430 nm, and emission was detected at 510 ± 10 nm, with an exposure time of 20 seconds. Fluorescence signals were acquired at 15-minute intervals. To calculate relative cell death intensity, fluorescence values at each time point were normalized to the corresponding intensity measured at time zero.

### Total protein extraction and analysis by BN-PAGE and SDS-PAGE Protein extraction

Four- to five-week-old *N. benthamiana nrc2/3/4* knockout plants were agroinfiltrated as described above in the “cell death assays” section with constructs of interest. Three days post-infiltration, six leaf discs were harvested per sample for protein analysis. Leaf tissue was homogenized using a Geno/Grinder tissue homogenizer. For SlNRC3 extractions, ground tissue was resuspended in GTMN buffer [10% [v/v] glycerol, 50 mM Tris-HCl (pH 7.5), 5 mM MgCl_2_, and 50 mM NaCl] supplemented with 10 mM dithiothreitol (DTT), 1× protease inhibitor cocktail (Sigma-Aldrich), and 0.2% [v/v] NP-40 substitute (Sigma-Aldrich). For NbNRC4, GHMN buffer [10% [v/v] glycerol, 50 mM HEPES (pH 7.5), 5 mM MgCl_2_, and 50 mM NaCl] supplemented with 10 mM DTT, 1× protease inhibitor cocktail, and 1% [w/v] digitonin (Sigma-Aldrich) was used. Samples were incubated on ice for 10 minutes with brief vortexing every 2 minutes to facilitate extraction. The lysates were then centrifuged at 5,000 ×g for 20 minutes at 4 °C. The resulting supernatants were collected and used for BN-PAGE and SDS-PAGE assays.

### BN-PAGE and SDS-PAGE analysis

For blue native polyacrylamide gel electrophoresis (BN-PAGE), clarified protein extracts were diluted according to the manufacturer’s instructions with NativePAGE™ 5% G-250 sample additive, 4× NativePAGE Sample Buffer, and deionized water. Samples were loaded onto 3%– 12% Bis-Tris NativePAGE™ gels (Invitrogen) and electrophoresed alongside SERVA Native Marker (SERVA). Following separation, proteins were transferred to polyvinylidene difluoride (PVDF) membranes using NuPAGE™ Transfer Buffer and the Trans-Blot Turbo Transfer System (Bio-Rad), according to the manufacturer’s protocol. Membranes were fixed in 8% acetic acid for 15 minutes, rinsed with water, and air-dried. To visualize native protein standards, membranes were reactivated with ethanol prior to immunoblotting.

For SDS-PAGE, protein samples were mixed with SDS loading buffer and denatured at 72 °C for 10 minutes. Following centrifugation at 5,000 ×g for 3 minutes, the supernatants were loaded onto 4%–20% Mini-PROTEAN^®^ TGX™ precast gels (Bio-Rad) and resolved alongside the PageRuler™ Plus Prestained Protein Ladder (Thermo Scientific). Proteins were transferred to PVDF membranes using the Trans-Blot Turbo Transfer System with the supplied transfer buffer, following standard procedures.

### Immunoblotting and detection

Membranes were blocked in 5% (w/v) non-fat dry milk in Tris-buffered saline containing 0.01% Tween-20 (TBS-T) for 1 hour at room temperature. Blots were then incubated overnight at 4 °C with horseradish peroxidase (HRP)-conjugated primary antibodies diluted 1:5,000 in 5% milk/TBS-T. The following antibodies were used: anti-GFP (B-2, Santa Cruz Biotechnology), anti-V5 (V5-10, Sigma-Aldrich), anti-FLAG (M2, Sigma-Aldrich), and anti-HA (clone 3F10, Roche).

Signal detection was performed using Pierce™ ECL Western Blotting Substrate (32106, Thermo Fisher Scientific), with up to 50% SuperSignal™ West Femto Maximum Sensitivity Substrate (34095, Thermo Fisher Scientific) added for enhanced sensitivity when needed. Chemiluminescence was imaged using either an ImageQuant LAS 4000 or an ImageQuant 800 system (GE Healthcare Life Sciences). Membrane loading controls were visualized by staining with Ponceau S (Sigma-Aldrich) or Ponceau 4R (AG Barr).

### Protein purification

Approximately 30 leaves from *Nicotiana benthamiana nrc2/3/4* knockout plants were agroinfiltrated as described above in the “cell death assays” section to transiently express SlNRC3^EEE^-3×FLAG, Rx-V5, PVX-CP-eGFP, and AVRcap1b-6×HA. Leaf tissue was harvested three days post-infiltration and immediately snap-frozen in liquid nitrogen. Samples were stored at –80 °C until protein extraction, which was completed on the same day for each preparation.

For extraction, frozen tissue was ground to a fine powder using a pre-chilled mortar and pestle under liquid nitrogen. A total of 20 g of ground tissue was resuspended in 80 mL of ice-cold extraction buffer (100 mM Tris-HCl, pH 7.5, 150 mM NaCl, 1 mM EDTA, 10% [v/v] glycerol, 10 mM DTT, 0.4% [v/v] IGEPAL, and one tablet of cOmplete™ EDTA-free protease inhibitor cocktail [Sigma]). After thorough vortexing and homogenization, the lysate was centrifuged at maximum speed for 20 minutes at 4 °C. The supernatant was then filtered through Miracloth (Merck) and transferred to a fresh tube and subjected to a second round of centrifugation under the same conditions. The resulting clarified extract was filtered through Miracloth (Merck) to remove residual debris.

The filtered extract was incubated with 200 μL of anti-FLAG M2 affinity gel (Sigma-Aldrich) at 4 °C for 90 minutes with continuous rotation to ensure uniform mixing. Following incubation, the protein-bound resin was collected on an open gravity-flow column and washed with 10 mL of wash buffer (100 mM Tris-HCl, pH 7.5, 150 mM NaCl, 1 mM EDTA, 10% [v/v] glycerol, and 0.1% [v/v] IGEPAL). Beads were then transferred to a microcentrifuge tube, and bound proteins were eluted in 100 μL of elution buffer (100 mM Tris-HCl, pH 7.5, 150 mM NaCl, 1 mM MgCl_2_, 3% [v/v] glycerol) supplemented with 0.5 mg/mL 3×FLAG peptide. Eluted proteins were analyzed by SDS-PAGE to assess purity and integrity. Purified proteins were subsequently used for cryo-electron microscopy (cryo-EM) analysis.

### Negative staining

Freshly prepared sample was diluted and applied to glow-discharged 300-mesh copper grids coated with continuous carbon (Agar Scientific) and stained with 2% (w/v) uranyl acetate in H_2_O. Grids were imaged manually on an FEI Talos F200C microscope operating at 200 KeV with a Gatan OneView camera (Gatan). Images (30 - 50 for each sample) were taken at a nominal magnification of 45,000 x corresponding to a pixel size of 2.87 Å. Micrographs were imported into CryoSPARC (v4.6.2) (*39*) where blob-picking was used to pick particles which were then subjected to 2D classification.

### Cryo-EM sample preparation and data collection

Sample preparation for all samples was identical. Quantifoil R 2/1 grids (Quantifoil Micro Tools GmbH) were coated with graphene oxide prior to plunge-freezing and used for both samples. Sample (∼3.5 μL, ∼0.3 mg/mL) was applied to glow-discharged grids and plunge-frozen into liquid ethane using a Leica GP2 Plunge Freezer (Leica Microsystems) operating at 4°C and a humidity of 90%. Grids were loaded into an FEI Titan Krios (Thermo Fisher Scientific) and screened for suitable particle density and ice quality. Movies were collected on the same microscope operating at 300 KeV with an energy filter inserted (slit width of 20 eV for SlNRC3 resistosome and 10 eV for SlNRC3-AVRcap1b complexes) on a K3 direct electron detector (Gatan) using the EPU software (Thermo Fisher Scientific). Movies were collected with a total dose of ∼50 e^−^/Å^2^ over 50 fractions at a nominal magnification of 105,000 X resulting in a pixel size of 0.828 Å. Defocus ranges used were –1.5 to –2.7 μm for the SlNRC3 resistosome and –0.6 to –2.0 μm for both SlNRC3-AVRcap1b complexes. A total of 6,150 movies were collected for the SlNRC3 resistosome, 7,475 movies from SlNRC3-AVRcap1b^WT^, and 7,555 movies for SlNRC3- AVRcap1b^P92E^. For all datasets, movies were collected with four exposures per hole. CryoSPARC (v4.6.2) was used for all processing.

#### SlNRC3 resistosome

Motion-corrected and CTF-estimated micrograph stacks were curated for suitable ice thickness and particle distribution. The resultant micrographs (6,118) were then used as input for blob picking to generate an initial particle stack. Particles were then extracted, binned 2x, then subjected to iterative rounds of 2D classification and template picking. Clear 2D classes that exhibited features of a hexameric assembly were then used to create an *ab-initio* reconstruction with C1 symmetry. The particles were then re-extracted to generate a stack of 234,243 unbinned particles which were used as input for a homogeneous refinement job with C6 symmetry and per-particle defocus and per-group CTF parameters optimized. Further improvement to the LRR regions of the map was achieved by using an 8 Å filtered map of the homogeneous refinement as input for non-uniform refinement. This produced the final consensus map at 2.85 Å.

#### SlNRC3-AVRcap1b complexes

Both SlNRC3-AVRcap1b^WT^ and SlNRC3-AVRcap1b^P92E^ datasets were processed in parallel for comparison. Motion-corrected and CTF-estimated micrograph stacks were curated for suitable ice thickness and particle distribution. The resultant micrographs (7,206 for AVRcap1b^WT^ and 7,363 for AVRcap1b^P92E^) were then used as input for blob picking to generate an initial particle stack. Particles were extracted, binned 2x, and then subjected to multiple rounds of 2D classification to generate clean classes. Initial 2D classification and *ab-initio* reconstructions yielded identical results for both complexes showing clear features of three SlNRC3 protomers bound by AVRcap1b. However, preferred orientation was clear in both datasets at this stage. Further refinement and 2D classification rounds were performed on both datasets, however the severe orientation bias of the SlNRC3-AVRcap1b^WT^ complex precluded generation of a high-resolution map.

For SlNRC3-AVRcap1b^P92E^ three *ab-initio* models with C1 symmetry were generated, which produced one class consisting of 304,153 particles (54% of all particles) with clear density for three SlNRC3 protomers and AVRcap1b. Initial refinement of this class revealed orientation bias that resulted in anisotropic density. Further 2D classification revealed two minor classes that were absent in the initial 2D classes. When pooled together, these particles produced an *ab-initio* class with more favorable orientation distribution. This particle stack was then further cleaned up by iterative 2D-classification and hetero-refinement to remove junk particles and contaminant hexameric particles. The final clean particle set was then re-extracted with a box size of 400 pixels resulting in 267,820 particles. Two rounds of non-uniform refinement with optimized per-particle defocus and CTF parameters produced a consensus reconstruction at a resolution of 3.01 Å. This consensus map showed clear density for a majority of the map, however the density for the N-terminus of AVRcap1b was comparatively weak. To improve the density of this region, a mask was created around this region and 3D-variability analysis was performed to identify particle clusters that contained better density for this region. This identified 149,587 particles showing clear density that were then used for local refinement with a Gaussian prior applied which produced a map with a resolution of 3.21 Å. These two maps were merged to create a composite map using the vop tool in ChimeraX (*40*). All reported resolutions use a Fourier shell correlation (FSC) cutoff of 0.143.

### Model building and refinement

#### SlNRC3 resistosome

AlphaFold 3 (*41*) was used to predict a model of the SlNRC3 resistosome by inputting 6 copies of the SlNRC3 protein lacking the first 18 amino acids. One protomer of this prediction was docked into the consensus EM map in ChimeraX and then manually inspected and refined in Coot (version 0.9.8.95) (*42*). Real-space refinement was carried out in Phenix (version 1.21.2) (*43*). This model was then further refined using ISOLDE (version 1.8) (*44*) within ChimeraX to fix Ramachandran and rotamer outliers and generally improve the quality of the model. This protomer was then imported into Phenix where it underwent real-space refinement before applying non-crystallographic symmetry (NCS) (using C6 symmetry) to produce the final hexamer model. The final structure was validated using cryo-EM validation and MolProbity (*45*) in Phenix.

#### SlNRC3-AVRcap1b complex

Three protomers of the SlNRC3 resistosome model and an AlphaFold 3 model of AVRcap1b were manually fit into the consensus EM map. Due to slight differences in angle between the AVRcap1b EM density and the AlphaFold model, AVRcap1b was split into two models, one consisting of LWY2 and the other consisting of LWY3-7. Each was manually fitted separately and then later merged. Initial building and refinement were carried out in Coot and Phenix using real space refinement. This initial model was then imported into ISOLDE and further refined. Following ISOLDE refinement, the model was then refined a final time in Phenix. C1 symmetry was used throughout refinement. The final structure was validated using cryo-EM validation and MolProbity in Phenix. Interacting residues and buried surface area were analyzed with the PISA server (*46*) and Contact (as part of the CCP4 package (*47*)) using a distance cutoff of 5 Å. Contact geometry was further assessed by using the ‘H-bonds’ tool in ChimeraX.

### Structure prediction

All structural predictions were run on the AlphaFold 3 webserver (*41*). Confidence scores for the SlNRC3-AVRcap1b^LWY3-7^ predicted model were ipTM = 0.54, and pTM = 0.62.

### Conservation analysis

Conservation of AVRcap1b and homologs was assessed using ConSurf server (*48*) using the 184 RXLR-LWY homologs from *Phytophthora* identified from a PSI-BLAST search of AVRcap1b against the NCBI non-redundant protein database described previously (Data S2 of (*22*)).

### Data visualization

All maps and models were visualized in ChimeraX v1.8-1.9.

