## Supplementary Information for "A plant pathogen effector blocks stepwise assembly of a helper NLR resistosome"

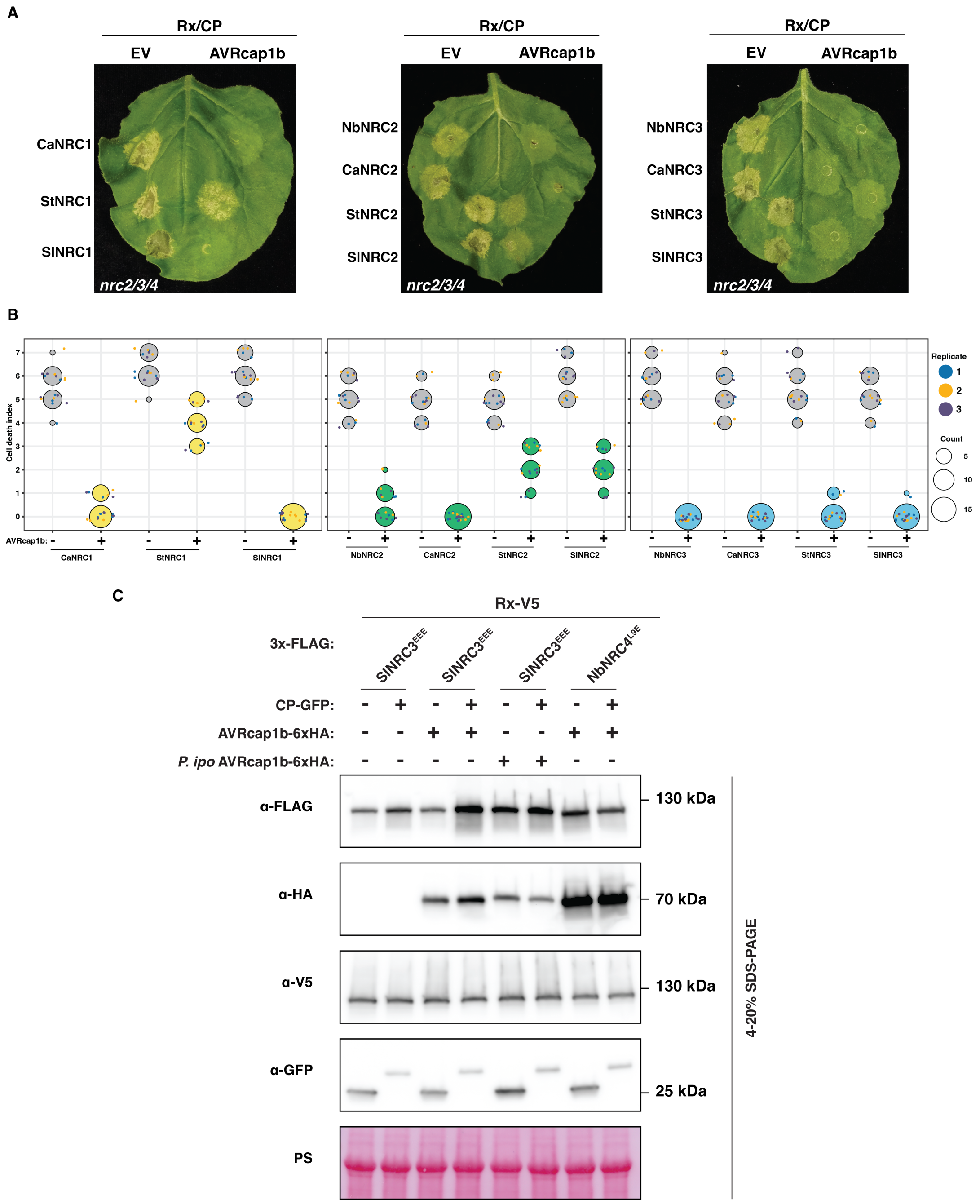


**Fig. S1. AVRcap1b differentially suppresses NRC1, NRC2 and NRC3 orthologs.** A) Representative *N. benthamiana nrc2/3/4* knockout leaves showing hypersensitive cell death following co-expression of the Rx sensor, *Potato virus X* (PVX) coat protein (CP), and various NRC1, NRC2 or NRC3 orthologs (Nb: *Nicotiana benthamiana*, Ca: *Capsicum annuum*, St: *Solanum tuberosum*, Sl: *Solanum lycopersicum*) together with either AVRcap1b or an empty vector (EV). B) Quantification of hypersensitive cell death intensity corresponding to the panel A. Cell death was scored on a modified 0–7 scale at 5 days post-infiltration. Scores are displayed as dot plots, where dot size reflects the number of replicates with identical scores. Data are representative of three independent biological replicates. C) SDS-PAGE analysis corresponding to the BN-PAGE experiment in **Fig. 1**, confirming protein accumulation of SlNRC3$^{EEE}$, NbNRC4$^{L9E}$, Rx, CP, and the indicated AVRcap1b variants. Total protein extracts were immunoblotted with the indicated antibodies (left). Approximate molecular weights (kDa) are shown on the right. Rubisco loading was visualized by Ponceau stain (PS).

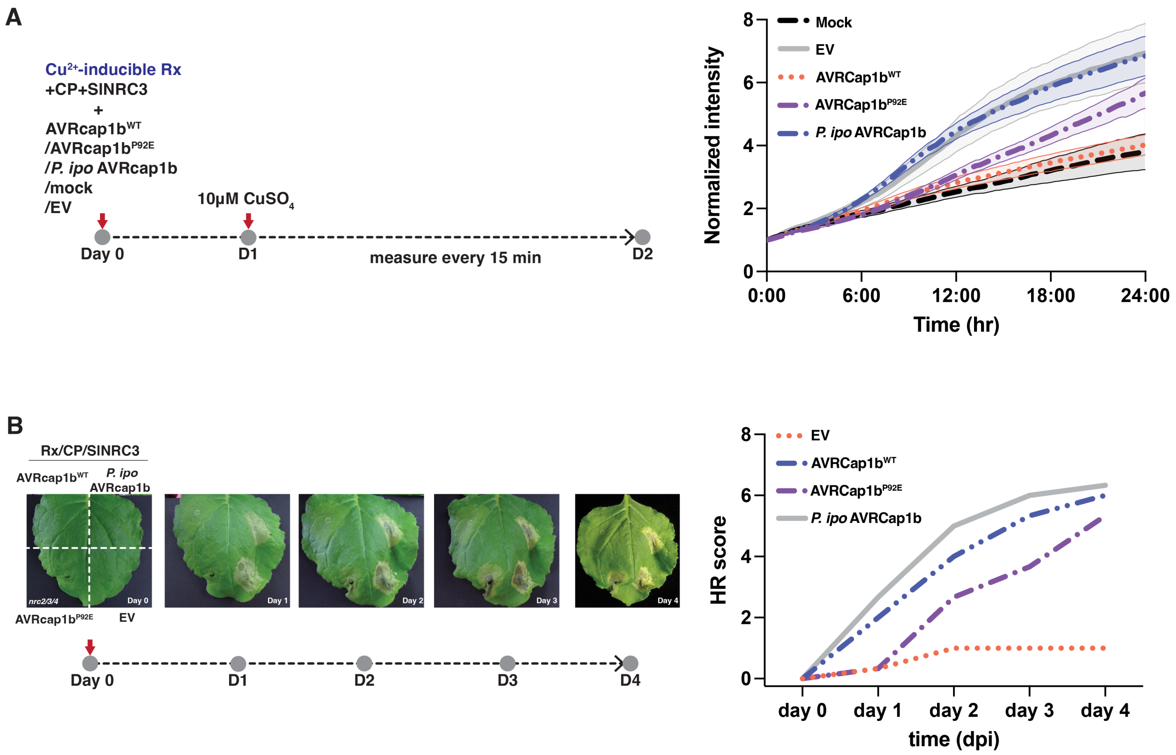


**Fig. S2. AVRcap1b partially suppresses immunity in the absence of TOL9a binding**. A) Time-course analysis of hypersensitive cell death using a copper inducible system. CP, SlNRC3, the AVRcap1b variants, or EV are infiltrated along with a copper inducible Rx construct. After 24 hours copper induction drives the expression of Rx and the autofluorescence (as a measure of cell death) is tracked for 24 hours (from day 1 to day 2). To calculate relative cell death intensity, fluorescence values at each time point were normalized to the corresponding intensity measured at time zero. Data represents 4 independent experiments with the mean and standard error of the mean (SEM) plotted. B) Time-course analysis of hypersensitive cell death following agroinfiltration of Rx/CP/SlNRC3 with AVRcap1b (either WT or P92E mutant which no longer binds TOL9a) with *P. ipomoeae* AVRcap1b and EV as controls. Data plotted is the mean of 2 technical replicates. Hypersensitive cell death was scored on a modified 0–7 scale at 5 days post-infiltration

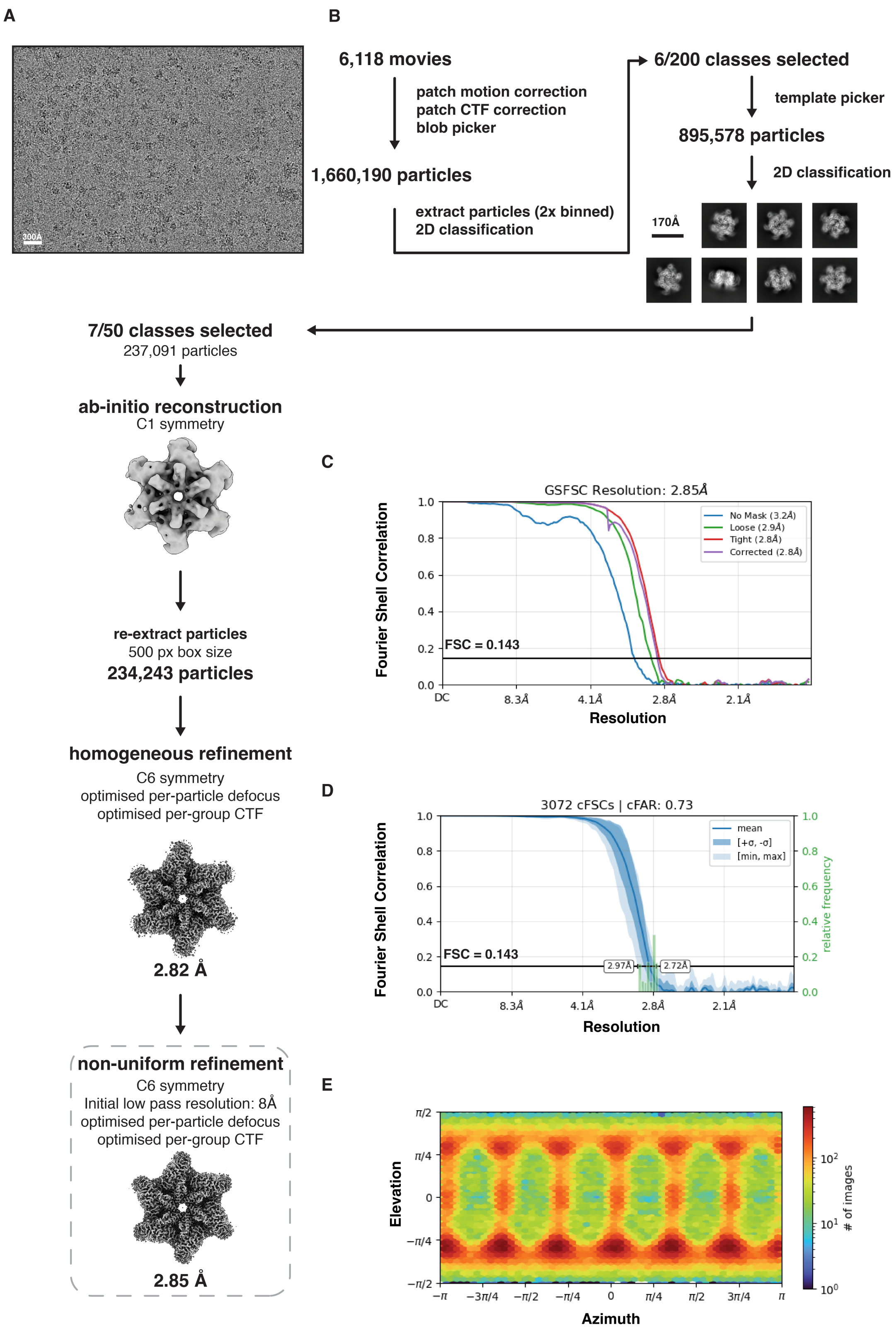


**Fig. S3. Cryo-EM data processing of the SlNRC3 hexamer.** A) Representative micrograph of the SlNRC3 hexamer. B) Flowchart of the SlNRC3 hexamer dataset showing 2D classes, *ab-initio* reconstructions and the final refined consensus map (outlined in a dotted box). Details in the methods section. C) Gold-standard Fourier shell correlation (FSC) curve of the final consensus reconstruction. D) 3D FSC curve of the consensus map. E) Angular distribution plot of the consensus map.

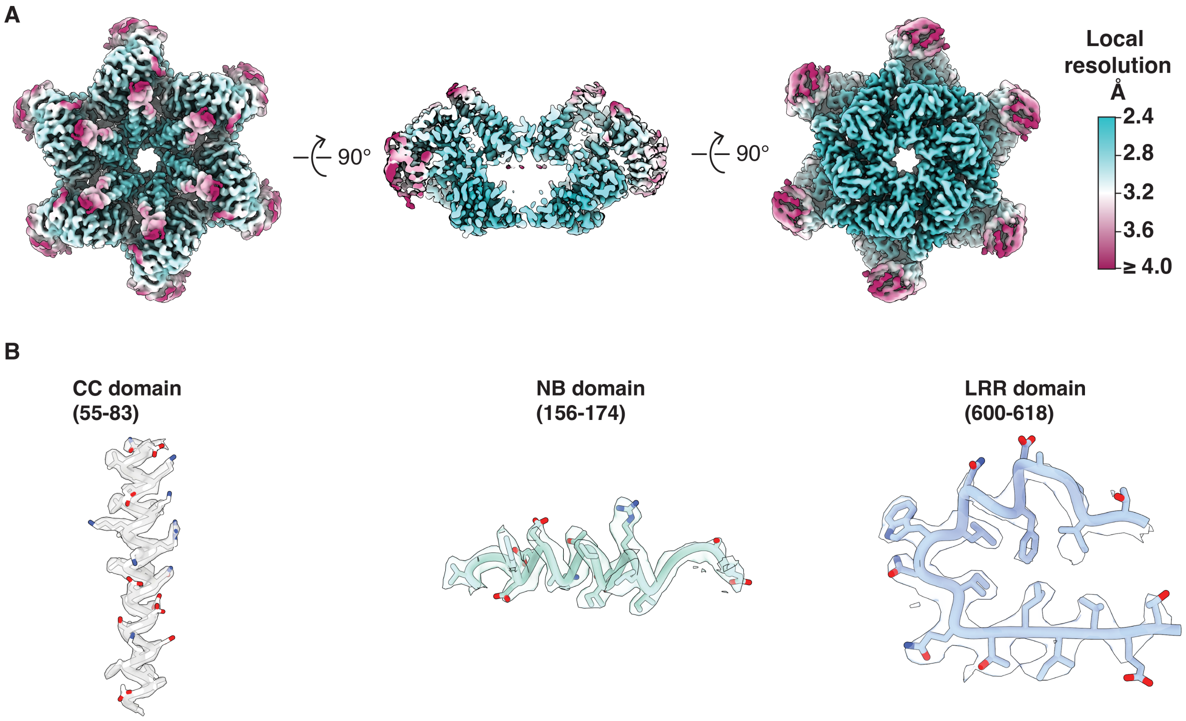


**Fig. S4. Quality assessment of SlNRC3 hexamer cryo-EM map.** A) Local resolution of the unsharpened SlNRC3 hexamer consensus map. B) Sharpened density around CC, NB and LRR regions of the map showing good map to model fit.

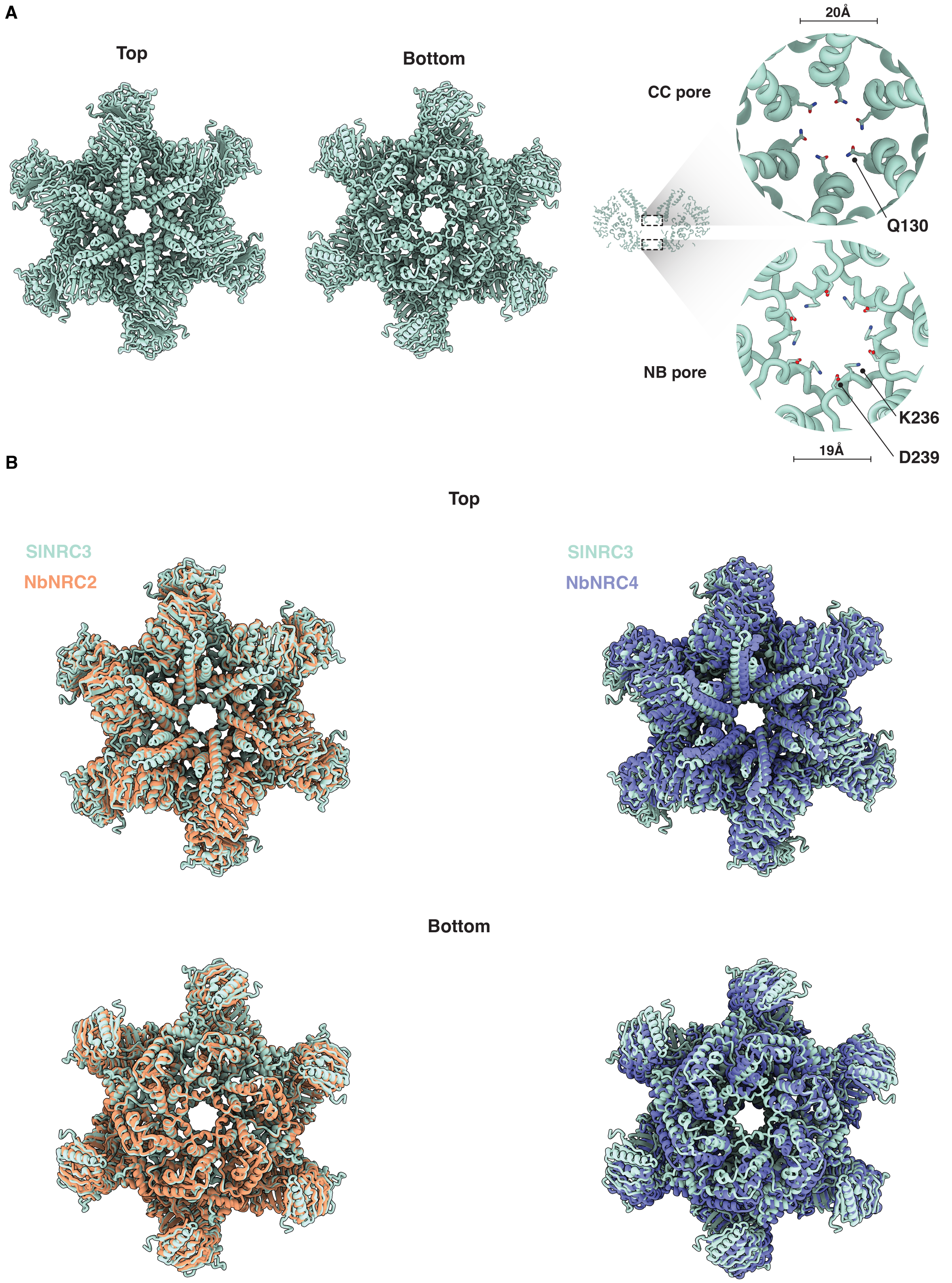


**Fig. S5. Activated SlNRC3 forms a resistosome that is comparable to NbNRC2 and NbNRC4 hexamers.** A) Overview of the atomic model of the SlNRC3 hexamer showing top and bottom views, as well as the CC- and NB-pores. Residues that form the pore constriction are depicted as sticks B) Superimposition of SlNRC3 hexamer with NbNRC2 (PDB: 9FP6) and NbNRC4 (PDB: 9CC8).

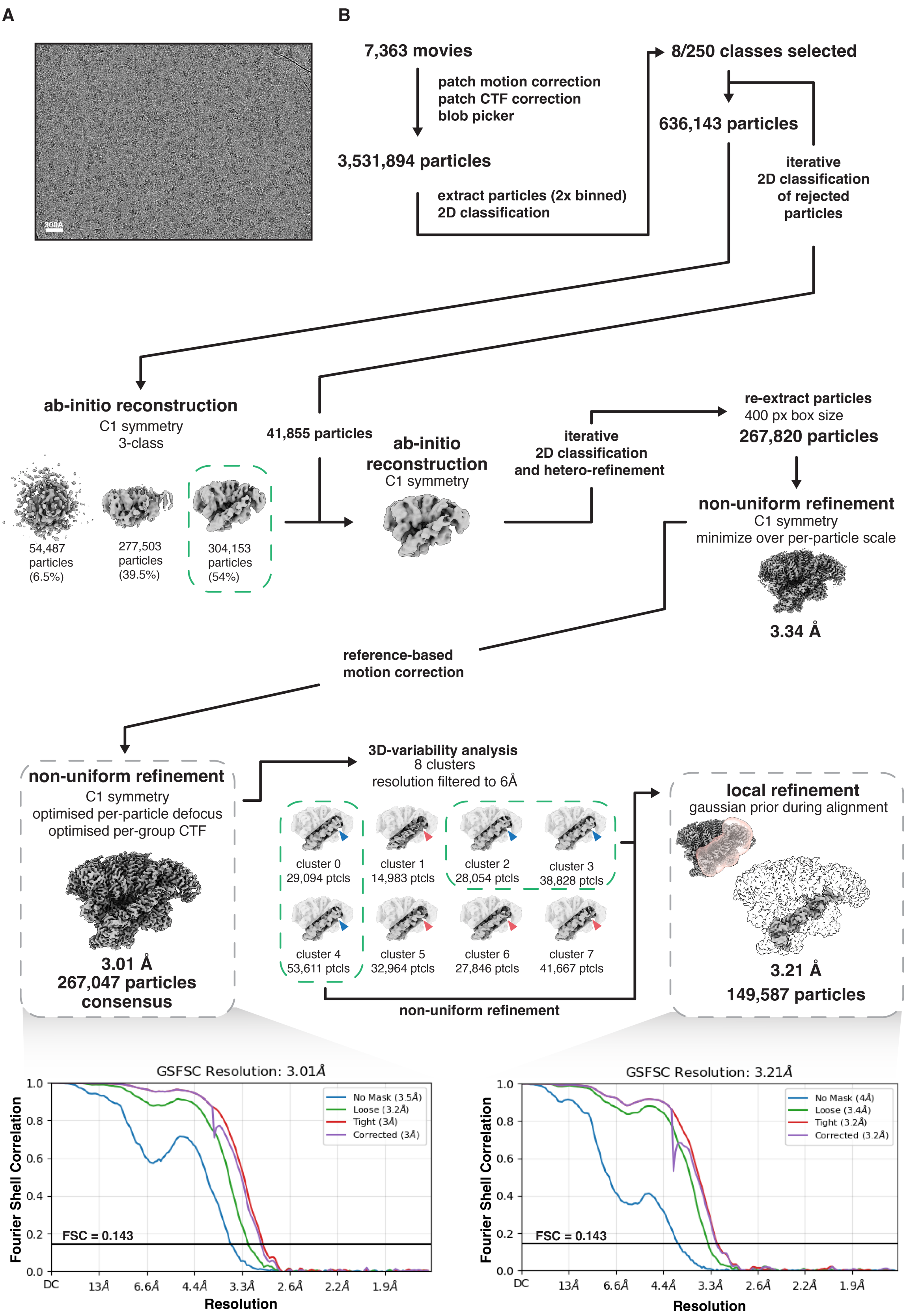


**Fig. S6. Cryo-EM data processing of the SlNRC3-AVRcap1b complex.** A) Representative micrograph of the SlNRC3-AVRcap1b complex. B) Flowchart of the SlNRC3-AVRcap1b dataset showing *ab-initio* reconstructions and the final refined maps (outlined in a dotted boxes) with associated FSC curves. Details in the methods section.

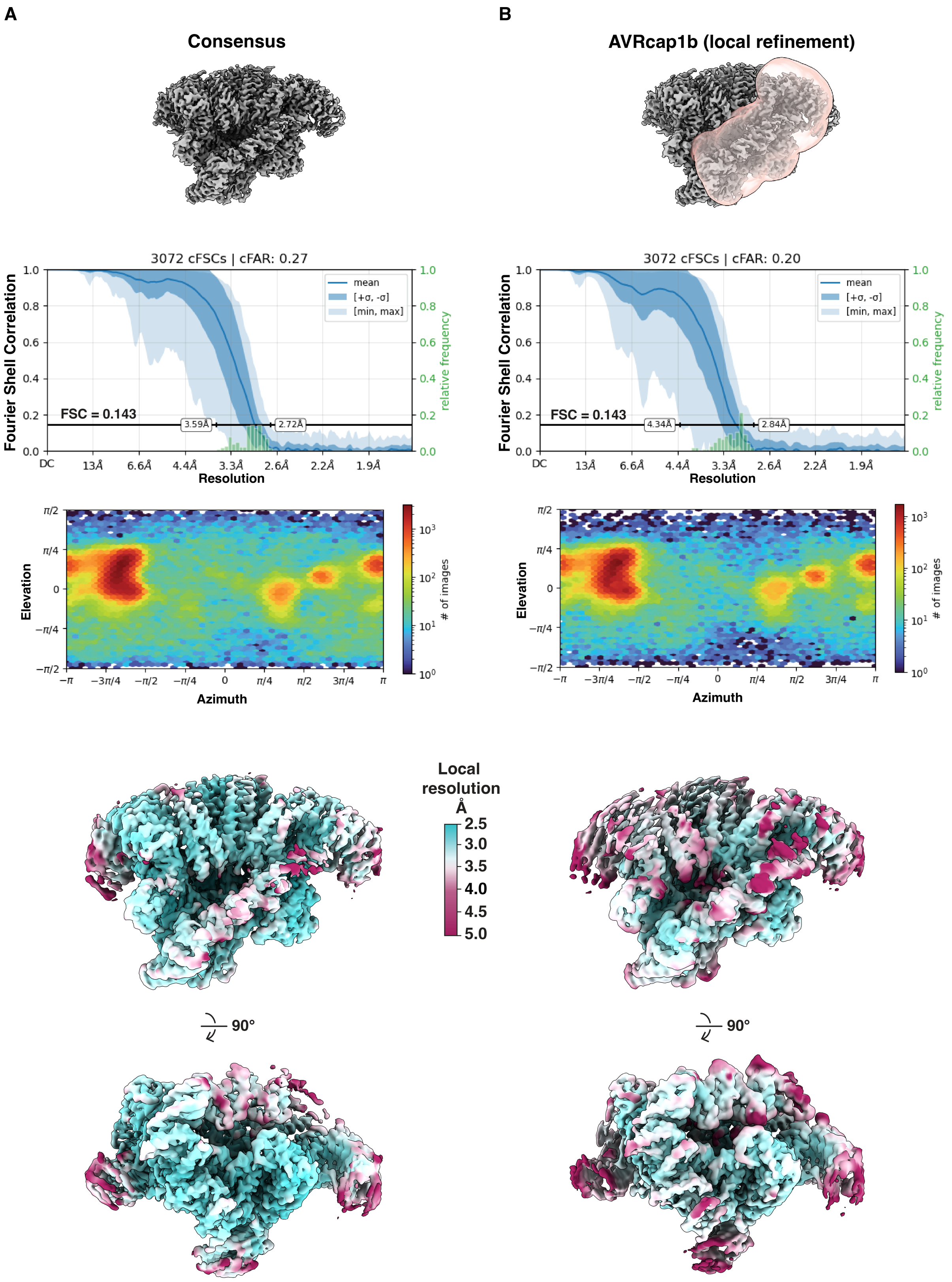


**Fig. S7. Quality assessment of SlNRC3-AVRcap1b complex cryo-EM maps.** A) 3D-FSC curve, angular distribution plot and local resolution assessment of the consensus reconstruction. B) 3D-FSC curve, angular distribution plot and local resolution assessment of the AVRcap1b locally refined map.

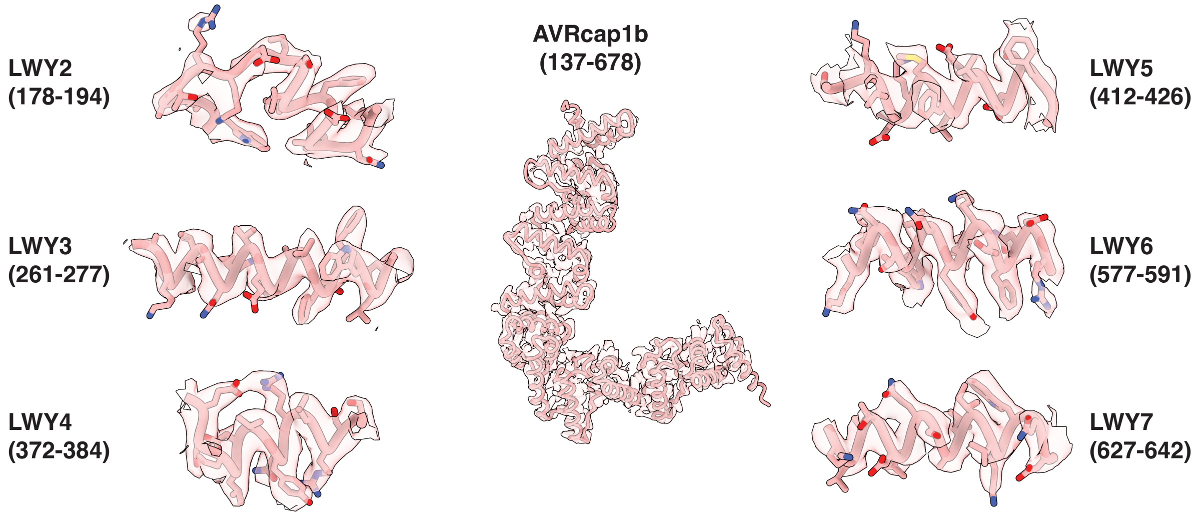


**Fig. S8. EM map quality assessment of AVRcap1b.** Composite map and fitted model of different regions of AVRcap1b.

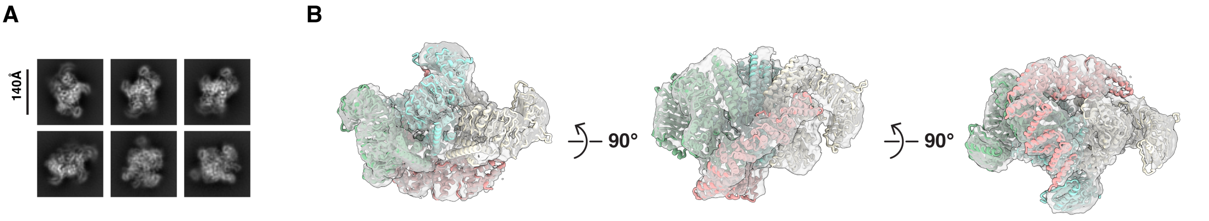


**Fig. S9. Cryo-EM analysis of SlNRC3 in complex with AVRcap1b$^{WT}$.** A) Representative 2D class averages. B) The model of SlNRC3-AVRcap1b$^{P92E}$ fits well into an *ab-initio* reconstruction of the SlNRC3-AVRcap1b$^{WT}$ complex.

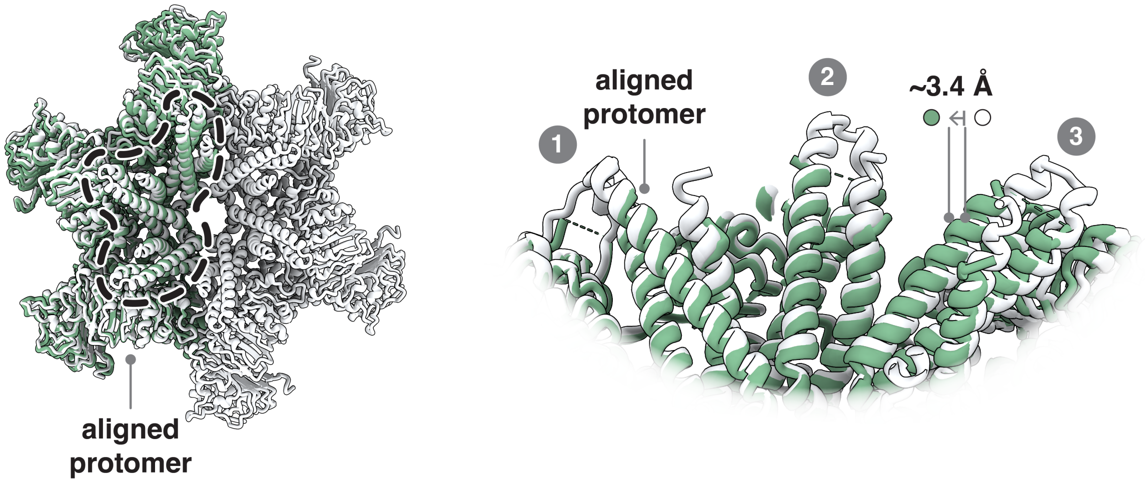


**Fig. S10. Comparison of AVRcap1b bound and unbound SlNRC3.** *Left*: Superimposition of SlNRC3-AVRcap1b intermediate complex (green) with the SlNRC3 hexameric resistosome (white). *Right*: Close-up of the dotted region showing minor deviation between the two structures. RMSD for the two aligned structures is 0.793 Å.

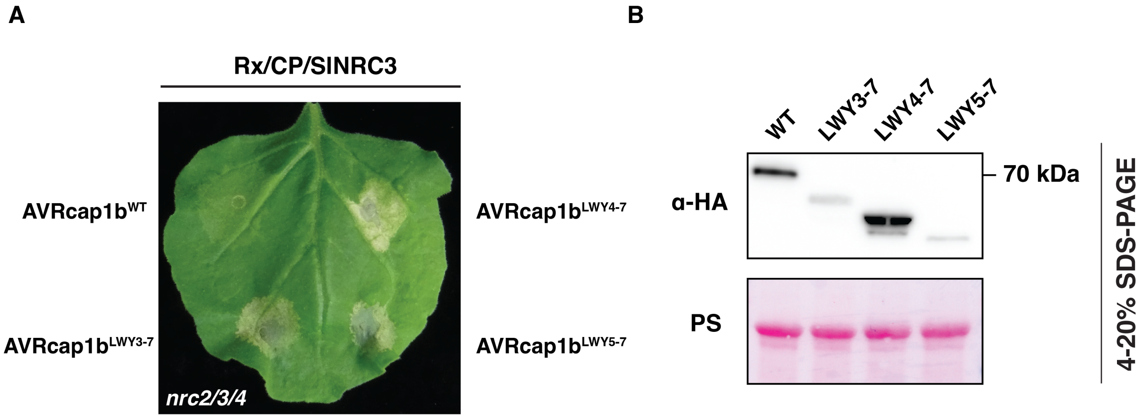


**Fig. S11. N-terminal truncations of AVRcap1b do not suppress SlNRC3-mediated cell death**. A) Representative *N. benthamiana nrc2/3/4* knockout leaf showing hypersensitive cell death response upon co-expression of the Rx sensor, CP, and SlNRC3$^{WT}$, together with full-length AVRcap1b or its N-terminal truncation variants. B) SDS-PAGE analysis confirming protein accumulation of the different AVRcap1b truncations. Total protein extracts were immunoblotted with the indicated antibody (left). Approximate molecular weights (kDa) are shown on the right. Rubisco loading was visualized by Ponceau stain (PS).

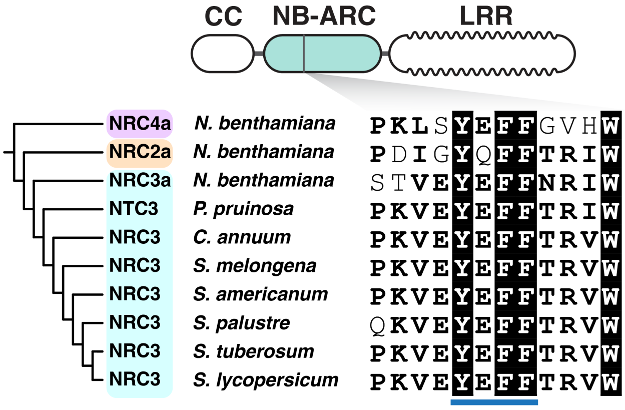


**Fig. S12. The 'YEFF' motif is conserved across NRC3 orthologs from Solanaceae species.** Cladogram and sequence alignment showing conservation of the YEFF motif across NRC3 orthologs from multiple plant species, along with NRC2 and NRC4 (indicated by a blue underline).

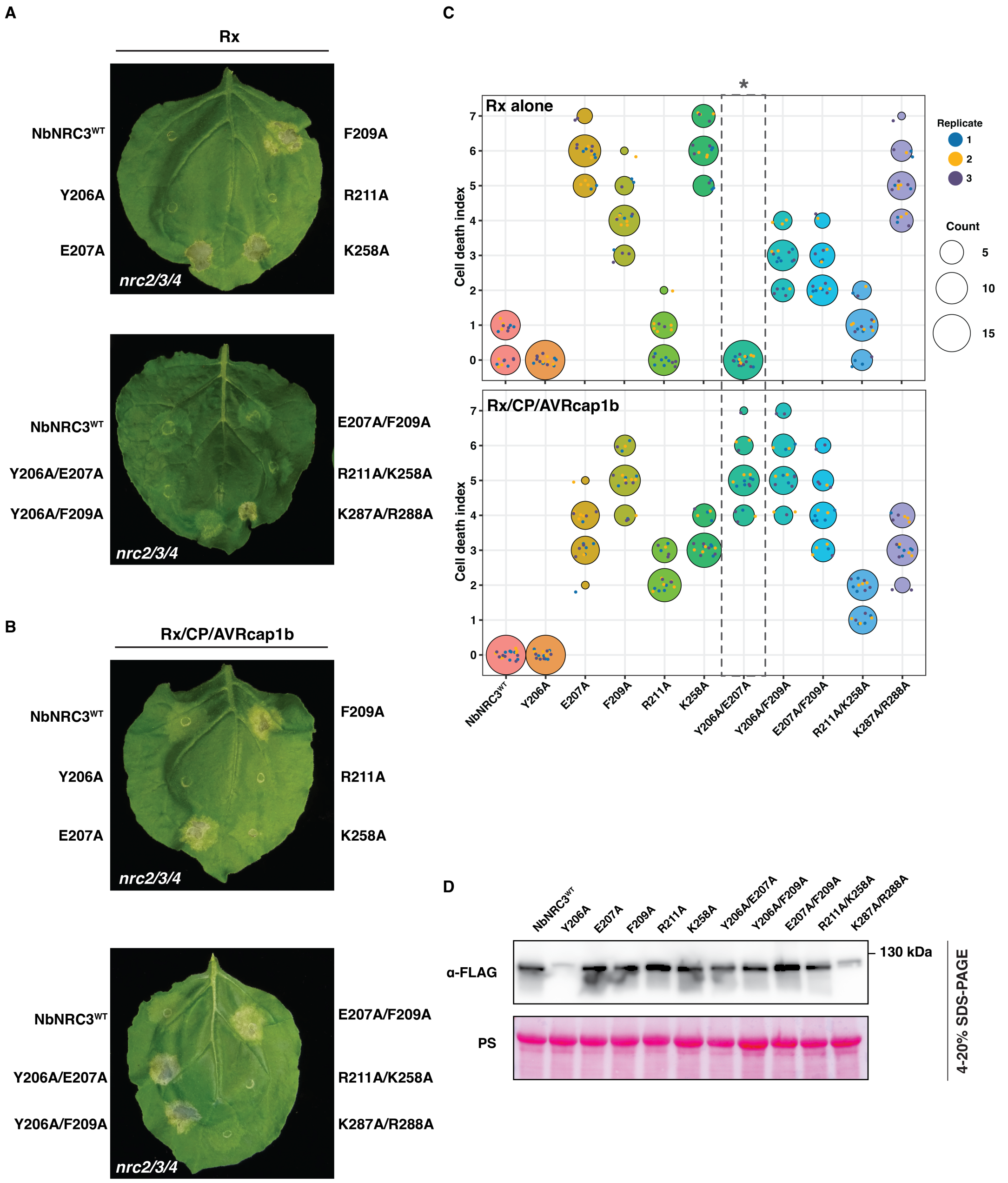


**Fig. S13. Several NbNRC3 mutants became insensitive to AVRcap1b suppression.** A) Representative *N. benthamiana nrc2/3/4* knockout leaves showing hypersensitive cell death after co-expression of NbNRC3$^{WT}$ or its mutants with Rx alone, revealing that a few mutants exhibit a trigger-happy phenotype. B) Representative *N. benthamiana nrc2/3/4* knockout leaves showing hypersensitive cell death following co-expression of the sensor Rx, CP, and either NbNRC3$^{WT}$ or its mutant variants, together with AVRcap1b. C) Quantification of hypersensitive cell death intensity corresponding to the panels (A, B). Hypersensitive cell death was scored on a modified 0–7 scale at 5 days post-infiltration. Scores are displayed as dot plots, where dot size reflects the number of replicates with identical scores. Data are representative of three independent biological replicates. A dotted box and asterisk indicate Y206A/E207A as a double mutant that is not trigger-happy and is not suppressed by AVRcap1b following Rx/CP activation. D) Immunoblots of total protein extracts from leaves expressing NbNRC3 variants, probed with the indicated antibody (left). Approximate molecular weights (kDa) of the proteins are shown on the right. Rubisco loading control was carried out using Ponceau stain (PS). The experiment was repeated three times with similar results.

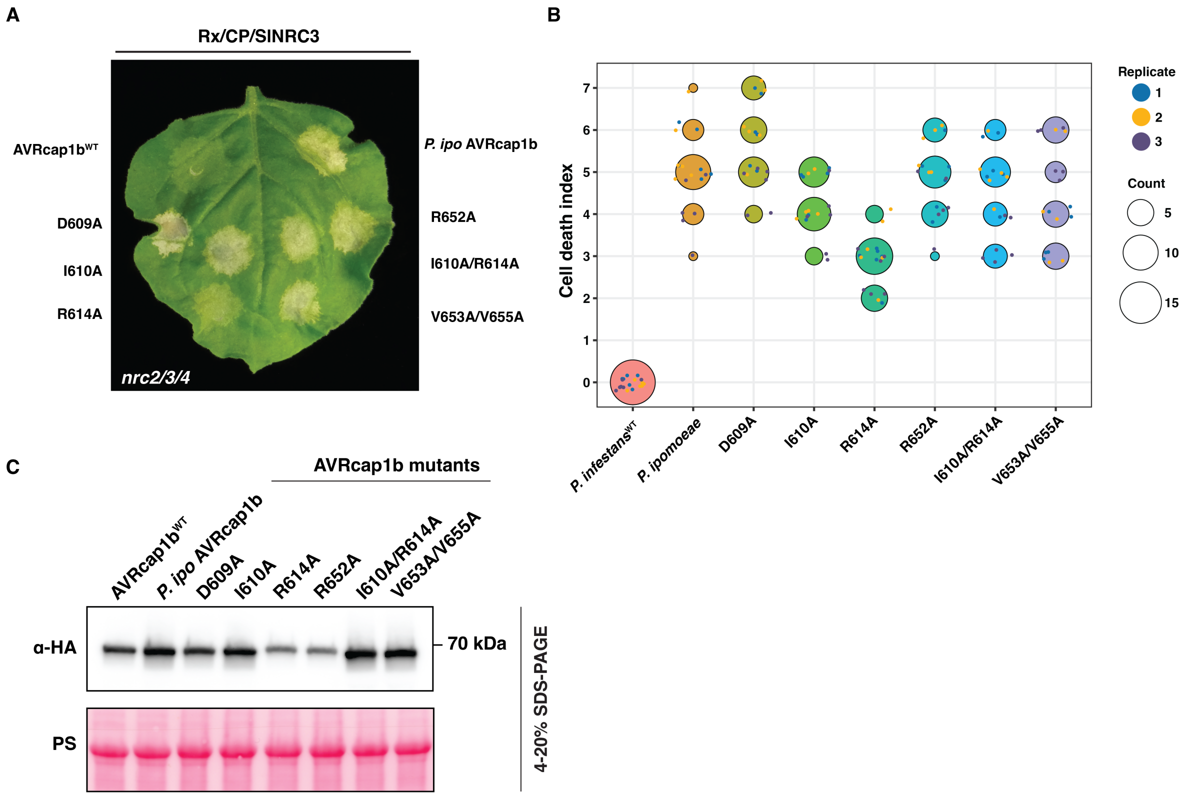


**Fig. S14. Mutations in the LWY7 module of AVRcap1b abolish its ability to suppress SlNRC3-mediated cell death.** A) Representative *N. benthamiana nrc2/3/4* knockout leaf showing hypersensitive cell death following co-expression of the Rx sensor, CP, and SlNRC3$^{WT}$, together with AVRcap1b or its LWY7 mutants. B) Quantification of hypersensitive cell death intensity corresponding to the panel A. Hypersensitive cell death was scored on a modified 0–7 scale at 5 days post-infiltration. Scores are displayed as dot plots, where dot size reflects the number of replicates with identical scores. Data are representative of three independent biological replicates. C) Immunoblots of total protein extracts from leaves expressing AVRcap1b LWY7 mutants, probed with the indicated antibody (left). Approximate molecular weights (kDa) of the proteins are shown on the right. Rubisco loading control was carried out using Ponceau stain (PS). The experiment was repeated three times with similar results.

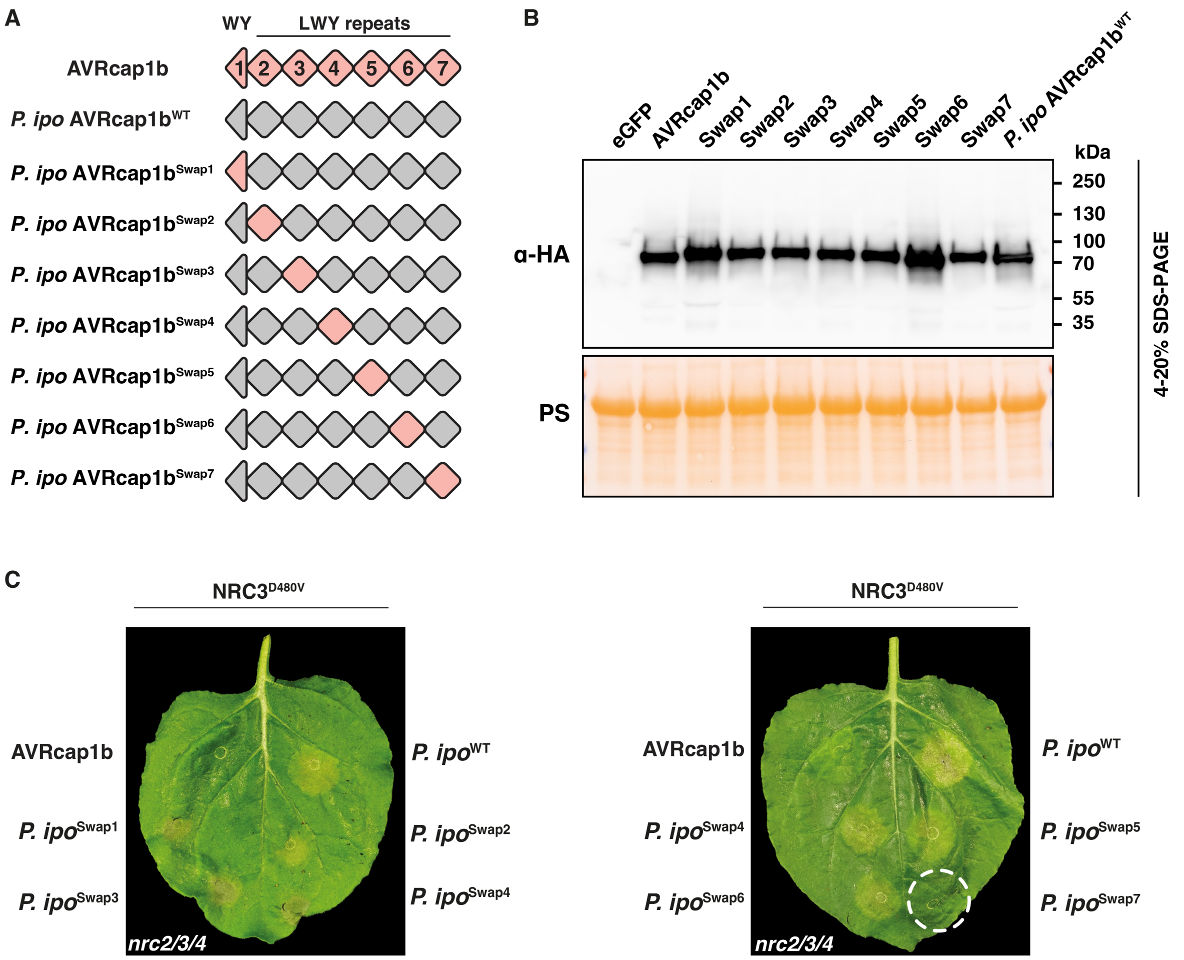


**Fig. S15. *P. ipomoeae* AVRcap1b gains the ability to suppress NRC3$^{D480V}$ upon swapping its LWY7 module with that of *P. infestans*.** A) Schematic representation of *P. ipomoeae* AVRcap1b chimeras generated by swapping individual WY/LWY modules from *P. infestans* AVRcap1b. B) Immunoblots of total protein extracts from leaves expressing AVRcap1b variants, probed with the indicated antibody (left). Approximate molecular weights (kDa) of the proteins are shown on the right. Rubisco loading control was carried out using Ponceau stain (PS). The experiment was repeated three times with similar results. C) Representative *N. benthamiana nrc2/3/4* knockout leaves showing hypersensitive cell death following co-expression of NRC3$^{D480V}$ together with AVRcap1b from *P. ipomoeae*, *P. infestans*, or their respective chimeras. The chimeric AVRcap1b from *P. ipomoeae* carrying the LWY7 module from *P. infestans* suppressed NRC3-mediated cell death (highlighted with a white dotted circle), underscoring the critical role of LWY7 in effector-mediated suppression.

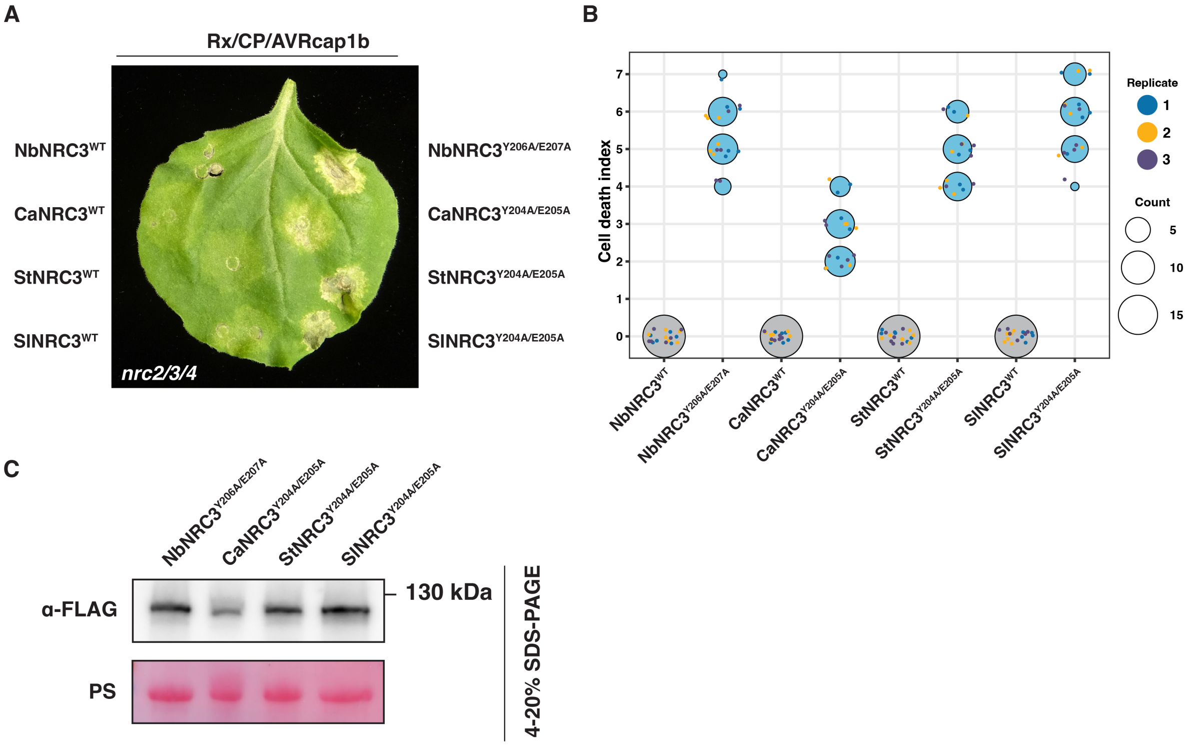


**Fig. S16. NRC3 orthologs with 'YEFF' motif mutations became insensitive to AVRcap1b suppression.** A) Representative *N. benthamiana nrc2/3/4* knockout leaf shows that introducing the equivalent Y206A/E207A double mutation into NRC3 orthologs (Nb: *Nicotiana benthamiana*, Ca: *Capsicum annuum*, St: *Solanum tuberosum*, Sl: *Solanum lycopersicum*) restores AVRcap1b-resistant immune responses. B) Quantification of hypersensitive cell death intensity corresponding to the panel A. Hypersensitive cell death was scored on a modified 0–7 scale at 5 days post-infiltration. Scores are displayed as dot plots, where dot size reflects the number of replicates with identical scores. Data are representative of three independent biological replicates. C) Immunoblots of total protein extracts from leaves expressing NRC3 ortholog YEFF mutants, probed with the indicated antibody (left). Approximate molecular weights (kDa) of the proteins are shown on the right. Rubisco loading control was carried out using Ponceau stain (PS).

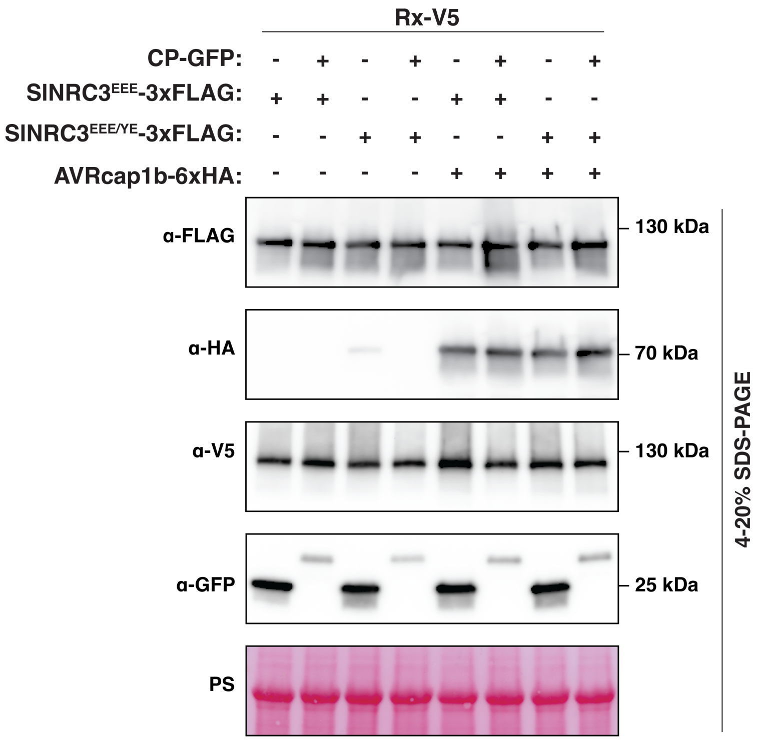


**Fig. S17. SDS-PAGE analysis corresponding to the BN-PAGE experiment in Fig. 5.** SDS-PAGE analysis corresponding to the BN-PAGE experiment in Fig. 5, confirming protein accumulation of SlNRC3$^{EEE}$, SlNRC3$^{EEE/YE}$, Rx, CP, and the indicated AVRcap1b variants. Total protein extracts were immunoblotted with the indicated antibodies and visualized by Ponceau stain (PS).

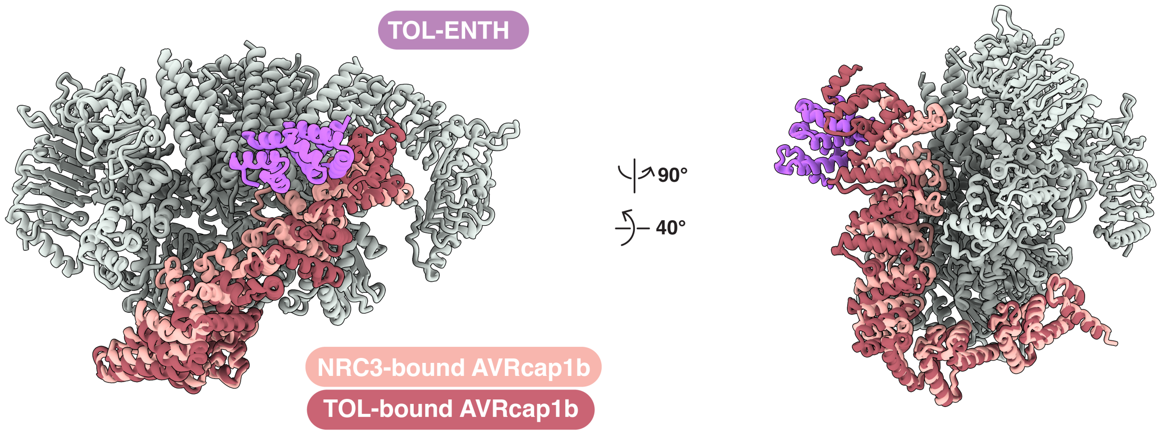


**Fig. S18. Comparison of the SlNRC3-AVRcap1b cryo-EM structure and AVRcap1b-NbTOL9a crystal structure**. Superimposition of the SlNRC3-AVRcap1b cryo-EM structure and the AVRcap1b-TOL9a crystal structure (PDB: 9RDC) shows that SlNRC3 and TOL9a bind to mutually exclusive interfaces of AVRcap1b, providing a hypothetical model of the ternary complex.

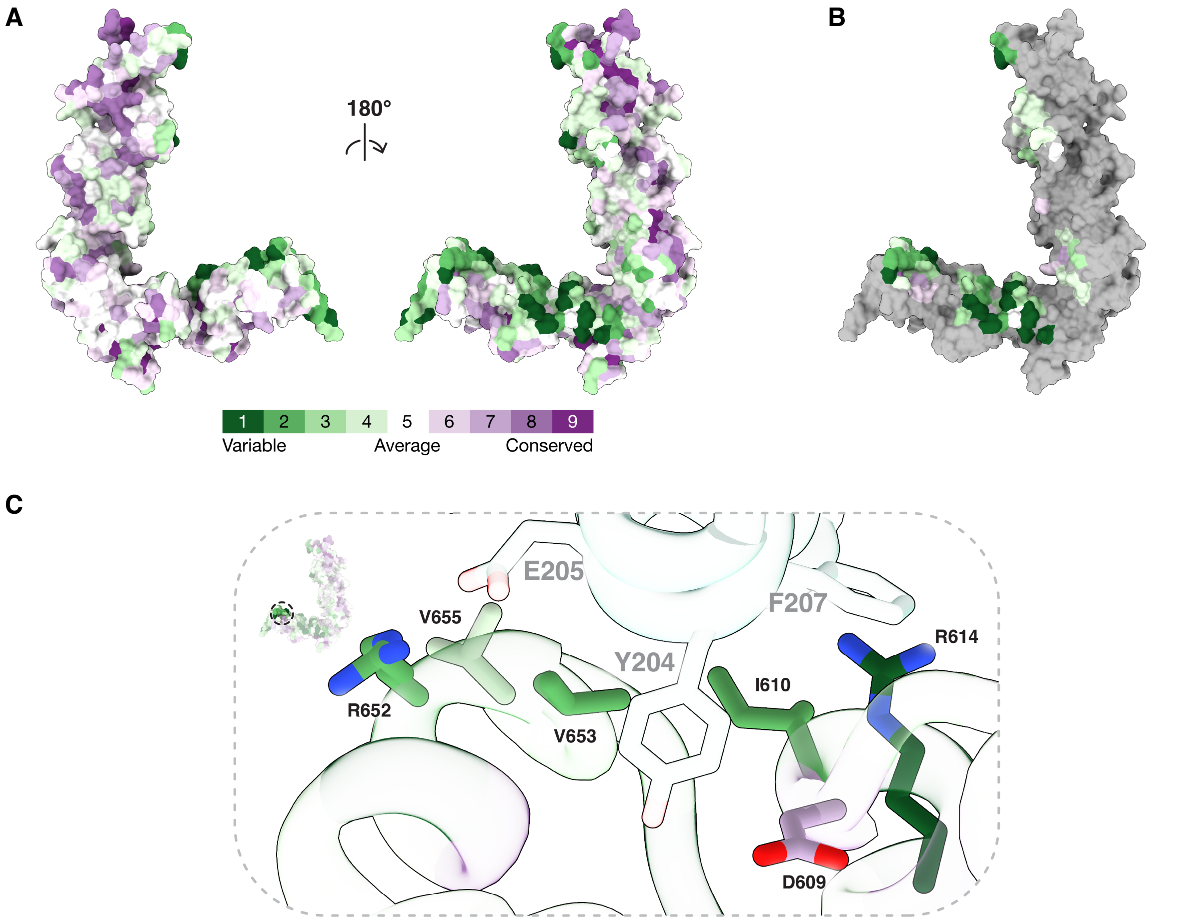


**Fig. S19. Conservation of AVRcap1b.** A) Surface representation of AVRcap1b from the SlNRC3-AVRcap1b structure showing sequence conservation of RXLR-LWY homologs from *Phytophthora* identified from a PSI-BLAST search of AVRcap1b against the NCBI non-redundant protein database and visualized using the ConSurf server. The left side shows the side of AVRcap1b not involved in SlNRC3 binding, and the right side shows the predominate face that contacts SlNRC3. B) Same as in A, but regions not involved in SlNRC3 binding are coloured in gray. C) Closeup view of the LWY7 region bound to the SlNRC3 YEFF motif. Residues that were mutated are shown as sticks and colored according to their level of conservation.

**Table S1: Cryo-EM data collection and refinement statistics**

| | SlNRC3 hexamer (PDB: 9RI9 EMDB: EMD-53990) | SlNRC3-AVRcap1b (Consensus) (EMDB: EMD-53988) | SlNRC3-AVRcap1b (Local refinement) (EMDB: EMD-53989) | SlNRC3-AVRcap1b (Composite map) (PDB: 9RIA EMDB: EMD-53991) |
|---|---|---|---|---|
| **Data collection and processing** | | | | |
| Magnification | 105,000 x | 105,000 x | | |
| Voltage (kV) | 300 | 300 | | |
| Electron exposure ($e^{-}$/Å$^2$) | 50.84 | 50.13 | | |
| Defocus range (µm) | -1.5 to -2.7 | -0.6 to -2.0 | | |
| Pixel size (Å) | 0.828 | 0.828 | | |
| Symmetry imposed | C6 | C1 | | |
| Initial micrographs | 6,150 | 7,555 | | |
| Final micrographs | 6,118 | 7,363 | | |
| Initial particle images | 1,660,190 | 3,531,889 | | |
| Final particle images | 234,243 | 267,047 | 149,587 | |
| Map resolution (Å) | 2.85 | 3.01 | 3.21 | |
| FSC threshold | 0.143 | 0.143 | 0.143 | |
| Sharpening *B* factor (Å$^2$) | -111.5 | | | -85.27 |
| **Model composition** | | | | |
| Non-hydrogen atoms | 41,892 | | | 24,742 |
| Protein residues | 5,208 | | | 3,075 |
| Ligand molecules | 6 | | | 3 |
| R.M.S deviations | | | | |
| Bond lengths (Å) | 0.004 | | | 0.004 |
| Bond angles (°) | 0.977 | | | 0.917 |
| **Model statistics and validation** | | | | |
| MolProbity score | 1.19 | | | 1.19 |
| Clashscore | 4.05 | | | 4.10 |
| Rotamer outliers (%) | 0.39 | | | 0.92 |
| Ramachandran plot | | | | |
| Favored (%) | 98.61 | | | 98.52 |
| Allowed (%) | 1.39 | | | 1.48 |
| Disallowed (%) | 0.00 | | | 0.00 |

### Table S2: SlNRC3-AVRcap1b table of contacts

| chain of 9RIA | Patch in Fig. 3B | SlNRC3 | | AVRcap1b | |
|---|---|---|---|---|---|
| | | | Gln 438 | | Ala 156 |
| | 1a | WHD | Gly 439 | | Leu 159 |
| | | | Pro 440 | | Tyr 160 |
| | | | | | Pro 161 |
| | | | | LWY2 | |
| | | | Tyr 40 | | |
| | | | Ala 43 | | Asn 193 |
| | | | Phe 44 | | Asn 195 |
| chain 'c' | | | Glu 47 | | Trp 196 |
| | | | Tyr 127 | | Gly 198 |
| | 1b | CC | Gln 130 | | |
| | | | Ala 131 | | Arg 236 |
| | | | Thr 133 | | Phe 237 |
| | | | Leu 134 | | Ser 238 |
| | | | Asp 135 | | Arg 239 |
| | | | Asp 136 | LWY3 | Lys 250 |
| | 1c | NB | Glu 243 | | Thr 258 |
| | 2a | CC | Leu 134 | | Arg 239 |
| | | | Ile 170 | | Arg 604 |
| | | | Lys 201 | | His 608 |
| | | | Glu 203 | | Asp 609 |
| | | | Tyr 204 | | Ile 610 |
| | | | Glu 205 | | Ala 611 |
| | | | Phe 206 | | Arg 614 |
| chain 'b' | | | Phe 207 | | Trp 640 |
| | 2b | NB | Arg 209 | LWY7 | Lys 645 |
| | | | | | Glu 649 |
| | | | Lys 256 | | Leu 650 |
| | | | Gly 257 | | Arg 652 |
| | | | Gly 258 | | Val 653 |
| | | | Lys 259 | | Phe 654 |
| | | | Gly 284 | | Val 655 |
| | | | Lys 285 | | Gly 656 |
| | | | Arg 286 | | Val 657 |

| chain of 9RIA | Patch in Fig. 3B | SlNRC3 | | AVRcap1b | |
|---|---|---|---|---|---|
| | 3a | CC | Gln 130 | LWY3 | Asp 241 |
| | | | Leu 134 | | Thr 295 |
| | | | Glu 143 | | Asp341 |
| | | | | | Tyr 385 |
| | | | Lys 146 | | Gly 386 |
| | | | | LWY4 | Thr 388 |
| | 3b | NB | Val 149 | | Asn 392 |
| | | | Val 150 | | Glu 393 |
| | | | Glu 152 | | Gln 395 |
| | | | Asp 153 | | Arg 429 |
| | | | Tyr 197 | LWY5 | |
| | | | Pro 200 | | Glu 466 |
| | | | Glu 203 | | Asp 479 |
| chain 'a' | | | Tyr 204 | | Asp 510 |
| | | | Glu 205 | | Arg 513 |
| | | | Phe 206 | | Ile 514 |
| | | | Phe 207 | | Ser 515 |
| | | | Thr 208 | | Gln 517 |
| | | | Arg 209 | | Pro 518 |
| | | | Val 210 | | Phe 519 |
| | 3c | NB | Ser 229 | | Arg 523 |
| | | | Lys 230 | | Tyr 545 |
| | | | Phe 231 | LWY6 | Tyr 548 |
| | | | Thr 232 | | Trp 549 |
| | | | Arg 233 | | Ile 552 |
| | | | Glu 252 | | Met 554 |
| | | | Gly 255 | | Glu 555 |
| | | | Lys 256 | | Thr 558 |
| | | | Gly 257 | | Arg 561 |
| | | | Gly 258 | | Ser 562 |
| | | | | | His 564 |
| | | | Lys 285 | | Asp 566 |
